# The Impact of AI-Based Modeling on the Accuracy of Protein Assembly Prediction: Insights from CASP15

**DOI:** 10.1101/2023.07.10.548341

**Authors:** Burcu Ozden, Andriy Kryshtafovych, Ezgi Karaca

**Affiliations:** Izmir Biomedicine and Genome Center, Izmir, Türkiye; Izmir International Biomedicine and Genome Institute, Dokuz Eylul University, Izmir, Türkiye; Protein Structure Prediction Center, Genome and Biomedical Sciences Facilities, University of California, Davis, California, USA

**Author notes:** **Correspondence** Ezgi Karaca.

**Keywords:** CASP, protein assembly, AF2-Multimer, quaternary structure prediction, protein-protein interaction, domain-domain interactions, deep-learning based modeling

## Abstract

In CASP15, 87 predictors submitted around 11,000 models on 41 assembly targets. The community demonstrated exceptional performance in overall fold and interface contact prediction, achieving an impressive success rate of 90% (compared to 31% in CASP14). This remarkable accomplishment is largely due to the incorporation of DeepMind’s AF2-Multimer approach into custom-built prediction pipelines. To evaluate the added value of participating methods, we compared the community models to the baseline AF2-Multimer predictor. In over 1/3 of cases the community models were superior to the baseline predictor. The main reasons for this improved performance were the use of custom-built multiple sequence alignments, optimized AF2-Multimer sampling, and the manual assembly of AF2-Multimer-built subcomplexes. The best three groups, in order, are Zheng, Venclovas and Wallner. Zheng and Venclovas reached a 73.2% success rate over all (41) cases, while Wallner attained 69.4% success rate over 36 cases. Nonetheless, challenges remain in predicting structures with weak evolutionary signals, such as nanobody-antigen, antibody-antigen, and viral complexes. Expectedly, modeling large complexes remains also challenging due to their high memory compute demands.

In addition to the assembly category, we assessed the accuracy of modeling interdomain interfaces in the tertiary structure prediction targets. Models on seven targets featuring 17 unique interfaces were analyzed. Best predictors achieved the 76.5% success rate, with the UM-TBM group being the leader. In the interdomain category, we observed that the predictors faced challenges, as in the case of the assembly category, when the evolutionary signal for a given domain pair was weak or the structure was large. Overall, CASP15 witnessed unprecedented improvement in interface modeling, reflecting the AI revolution seen in CASP14.

## INTRODUCTION

The understanding of protein interactions at the atomistic scale is crucial for studying cellular function. Experimental techniques like X-ray diffraction, NMR spectroscopy, and cryo-electron microscopy (cryo-EM) provide high-resolution structures of protein complexes^1^. However, in cases where experimental approaches face limitations, modeling becomes valuable^2–5^. In particular, homology modeling is employed when there is a resolved structure of an evolutionarily related complex, while docking is preferred in the absence of such a template. Various strategies, including coevolution integration and the use of available experimental data, have been implemented to enhance docking accuracy^6–12^. For intricate cases involving intertwined complexes, fold-and-dock strategies are employed^13,14^.

The CAPRI blind docking competition has been evaluating the state-of-the-art in assembly modeling since 2002 ^15,16^. In 2014, CAPRI joined forces with CASP to assess the prediction of protein complexes on a larger scale. Several rounds of CASP-CAPRI experiments have been conducted, shedding light on the capabilities and limitations of assembly modeling approaches^15,17–24^. A major limitation in protein complex modeling has been the absence of reliable templates for modeling the monomer structures of an assembly^20,25^. This limitation has been alleviated to a large extent with the release of AlphaFold2 (AF2), an artificial intelligence (AI) tool that has made unprecedented progress in tertiary structure prediction^26^. In CASP14, AF2 demonstrated high accuracy in modeling tertiary structure targets regardless of the prediction difficulty^26,27^. Consequently, the release of AF2 Protein Structure Database in 2022, with over 214 million predicted protein structures covering nearly all UniProt sequences, has significantly impacted the field of structural biology^28,29^.

Since the release of AF2, scientists have sought to incorporate this framework into their modeling pipelines. The simplest way to employ tertiary structure modeling methods to quaternary structure modeling was to join individual sequences of complex subunits into a longer, artificial sequence by means of adding an artificial glycine linker between monomers or introducing a sequence gap between multiple chains^30–33^. These approaches showed improvement over classical docking methodologies^34–36^. In 2021, DeepMind released AlphaFold-Multimer (AF2M, version 2.2), the multimeric version of AF2 specifically retrained on biological interfaces^33^. AF2M outperformed previously outlined AF2-monomer modifications in the case of heteromers. Although DeepMind did not participate in CASP15, they did so indirectly since the vast majority of assembly groups adopted AF2M in their modeling pipelines^37^.

In CASP15, to enhance the model accuracy, most of the groups experimented either with the multiple sequence alignment (MSA) input or with the AF2M sampling parameters. Among these, many benefitted from feeding their improved custom MSA into AF2M. A few utilized increasing the number of recycles (how many times the solutions are fed back to the AI-model), generating more models by starting from different random seeds or activating the dropout layers in the network. While tracing the specific AF2M modification strategies utilized by the community, we analyzed 41 assembly targets, considering the capabilities and limitations of the presented AF2M modifications. Our analysis shows where the community could push the boundaries of AI-based assembly modeling forward. It also highlights the remaining open challenges in the field.

## METHODS

### Assembly Assessment: General Concepts

Predictors in the CASP15 assembly category were provided with the stoichiometry information. We verified the symmetries of the targets in collaboration with the CAPRI Team by using AnAnaS^38^ and biological assemblies by using EPPIC^39^ and PISA^40,41^. Additionally, we the assessed interface properties, where needed by using PISA. Four targets, T119o, T1176o, T1176ov1, and T1184o, were canceled due to mismatch between the biological interfaces assigned by us and by the authors (Figure S1A). Targets H1171 and H1172 were multiple-conformation complexes spanning 1.5Å all-atom RMSD range. While assessing H1171 and H1172, we compared submitted models to all available conformers (two for H1171 and four for H1172) and reported the results for the top scoring model of each group.

Targets were classified into three prediction difficulty categories: *easy* if there is a structural template for the entire assembly, *medium* if there is a partial template for subunits or their interactions, and *hard* if there is no template. HHPred (https://toolkit.tuebingen.mpg.de/tools/hhpred)^42,43^ was used for template search. Additionally, we grouped targets based on their complex types, including homomeric (consisting of identical subunits), heteromeric (formed by different subunits), intertwined (with small segments or domains exchanged between subunits), nanobody-antigen, and antibody-antigen.

### Evaluation unit definitions

Evaluation of assembly targets requires establishing a structural correspondence between the constituent chains of the model and experimental structures. This proved to be a challenging and time-consuming task for complexes with many subunits. Recently, Gabriel Studer suggested a greedy heuristic approach (https://git.scicore.unibas.ch/schwede/casp15_ema) to solve the chain mapping problem^44^. This development allowed us to proceed with the evaluation of large complexes directly, without splitting them in subcomplexes as we did in CASP14^20^. However, we still needed to alter four targets.

*1) H1114* is an A4B8C8 bacterial assembly with C4 symmetry. In this complex, monomer A is forming an intertwined tetrameric tube, which interacts with monomer B. The prediction challenge in this complex was to form the intertwined tube and its interactions. To specifically focus on these interactions, we evaluated A4B2 assembly as a separate target *H1114v2* (Figure S1B). *2-4) H1166, H1167, H1168* are A1B1C1 immune complexes consisting of two antibody subunits (heavy and light chains) and an antigen molecule. To focus on the antibody-antigen interactions, we merged two antibody chains into a single chain, and formally assessed the complexes as hetero-dimers *H1166v1, H1167v1, H1168v1* (Figure S1C).

### Scoring and ranking

Submitted models were scored according to four metrics: two evaluating the accuracy of reproducing interfaces -Interface Contact Score (ICS, calculated as F1 score)^18^ and Interface Patch Score (IPS, calculated as Jaccard coefficient)^18^, and the other two evaluating overall model fold - Template Modeling (TM) score^45^, and Local Distance Difference Test (lDDT) score^46^.

ICS measures the relationship between the precision (P) and recall (R) of predicted inter-chain contacts. P is the fraction of the correct inter-chain contacts among all model inter-chain contacts. R is the fraction of correctly reproduced native inter-chain contacts. M and T represent the model and target. Two residues are considered in contact when at least two of their non-hydrogen atoms (one from each residue) are within 5Å from each other.

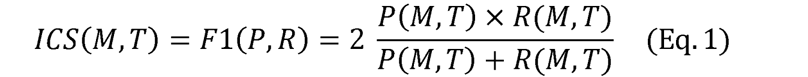

IPS is calculated as a Jaccard coefficient (J_c_) over the interface amino acids (I) predicted by the model (I_M_) compared to the target (I_T_):

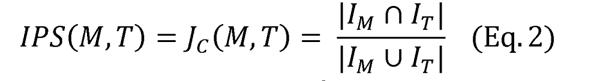

TM score measures the topological similarity of the model with respect to the target, where L and L_ali_ represent the target’s length and the number of equivalent residues in two structures, respectively. d_i_ is the distance between the i^th^ pair of equivalent residues in the two structures and score is normalized by scaling with d_0_.

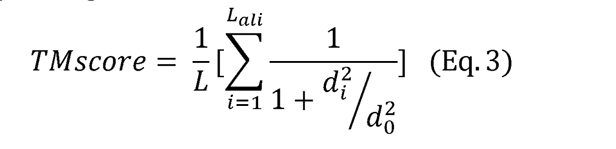

lDDT measures the difference in the interatomic distances within a model and the target. It is calculated by examining distances between pairs of atoms in the target that fall within a predefined threshold. If the difference in these distances is within a threshold (15 Å), the interaction is considered preserved in the prediction. The final score is determined by averaging the fraction of accurately modeled interactions.

We consider a model successful if it scores above 0.50 on all metrics. Focusing our analysis on the regions where complex subunits interact, we define three interface prediction accuracy zones: poor (ICS < 0.50), good (0.50 ≤ ICS < 0.75) and high (ICS ≥ 0.75). To investigate the interface prediction improvement across different CASP rounds, we carried out a comparative distribution analysis. For this, we collected the best ICS values generated for a given difficulty class during every round. The distributions of these values were then represented with box-and-whisker plots.

Participants were ranked based on their relative performance according to all four evaluation measures. A per-measure relative performance was calculated in terms of Z-scores reporting the number of standard deviations a particular model’s score is above or below the per-target population mean:

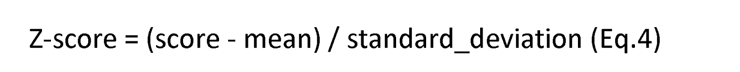

Z-scores were calculated for two different model sets: all submitted models or models designated as model_1 (i.e., top ranked models according to the predictor). Initially, Z-scores are computed on full model sets. Then outliers (models with Z-scores < -2) are removed, and Z-scores are re-calculated on the remaining models. Negative Z-scores are reset to 0 to avoid strong penalization of groups who underperformed. The final ranking score of a group is computed as a sum of combined per-measure Z-scores over all predicted targets:

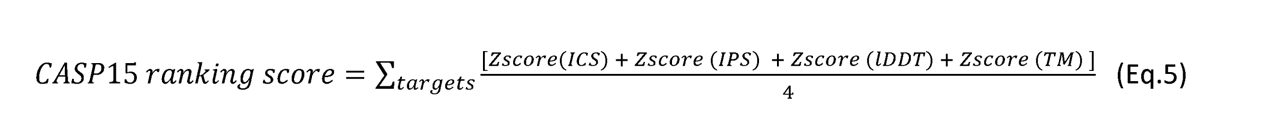

### Ranking on first models is considered the ‘official’ CASP ranking

The robustness of ranking is verified by the jackknife resampling. For this, initially, one target was excluded from the analysis, and the sum of Z-scores was calculated over the remaining targets. This process was repeated iteratively, excluding a different target at each step, until all targets were excluded once. For the final ranking, we utilized the mean scores obtained from the jackknifing protocol. The standard deviations from the jackknifed data set are used as a measure of the results’ variability.

### Naïve (baseline predictor)

Vast majority of CASP15 groups used the AF2M method as a starting point for their modeling. To help predictors with obtaining starting models, CASP organizers made an arrangement with Claudio Mirabello from the Elofsson group to run the public AF2M (version 2.2) method on all CASP15 targets and submit obtained models to CASP as predictions from the NBIS-AF2-Multimer group (NBIS-AF2M). In our evaluation, these predictions were used as the baseline models. Hereafter, NBIS-AF2M will be referred to as the baseline or AF2M^47^. We also used the underlying MSAs (deposited in http://duffman.it.liu.se/casp15/)^48^ to calculate the depth of sequence profiles. By default, the input MSAs were generated by JackHMMER v3.3^49^.The informational content of an MSA was evaluated with Neff, the number of effective sequences^50^. Neff is calculated as

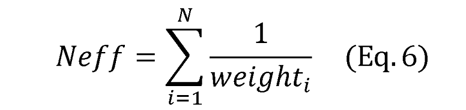

where N is the number of sequences in the MSA. For each sequence i, weight_i_ is the sum of weight_ij_ over all the sequences in the MSA, while weight_ij_ is the sequence identity between any homologues sequence i and j in the MSA. The minimum sequence identity between i and j was set to 80% identical residues.

For the targets with subunits, we calculated the multimeric Neff as

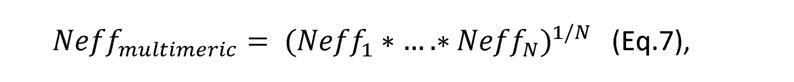

where N is the number of different subunits.

The baseline AF2M method was run with default parameters (no_recycles=3, no_seeds=5)^47^, except for T1173, T1174, and T1181, where NBIS team set the number of recycles to 25. Models for large targets -H1111, H1114, H1115, H1135, H1137 - were built by MolPC^51^. The final ranking was performed according to the model confidence score (0.8 · ipTM + 0.2 · pTM)^33^, where pTM is the predicted TM and ipTM is the same metric calculated over the interface.

To quantify the difference between the community performance and the baseline, we introduced a ΔICS score (Eq. 8). For each target, this score measures a gain in the ICS of the best community model over the AF2M one:

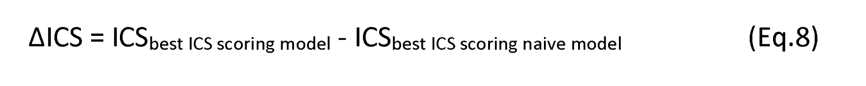

If needed, buried surface area (BSA) and noncovalent intermonomer interactions were calculated with the COCOMAPS tool (https://www.molnac.unisa.it/BioTools/cocomaps/)^52^.

### Interdomain Assessment: General Concepts

All multi-domain targets were considered as candidates for the interdomain assessment. If a tertiary structure target had multiple different interfaces, each of them was considered as a potential interdomain target^18^. Domain pairs with weak interfaces were disregarded, and only interfaces with ten or more interacting residue pairs were considered. For the evaluation of interdomain interactions we employed two interface metrics used in the assembly assessment: Interface Contact Score (ICS) and Interface Patch Score (IPS), and, additionally, the global Quaternary Structure (QSglobal) score^53,54^. The predictors were ranked according to the cumulative Z-score function:

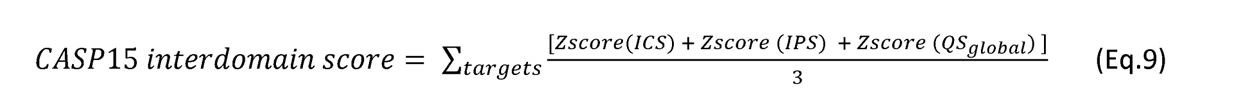

Z-scores for each metric were calculated on first models using the same procedure as explained above for ranking assembly groups.

### Visualization software

All plots in this paper were generated by using Python 3.8.10 and numpy, seaborn, matplotlib, pandas, and statistics libraries^55–59^.

The structure figures were generated with PyMOL(TM) 2.5.2^60^.

## RESULTS AND DISCUSSION

### General Overview of the CASP15 Assembly Prediction Category

In CASP15, 40 protein complexes were offered as prediction targets (Table 1, Figure S2). 29 of them were solved by X-ray crystallography, and the remaining 11 by cryo-EM. Half of the structures exhibited various degrees of cyclic symmetries. The target complexes contained from two to 27 subunits, with dimers prevailing. The homomeric cases comprised half of the assembly targets, including ten intertwined assemblies. The heteromeric cases consisted of three intertwined complexes (H1114, H1114v2 and H1137), one host-pathogen complex (H1129), and eight immunity complexes - five nanobody-antigen (H1140-H1144) and three antibody-antigen (H1166-H1168). The nanobody-antigen series involved binding of five different nanobodies to the catalytic domain of mouse CNPase, an enzyme responsible for regulating cyclic nucleotide synthesis^61^. While sharing a moderate sequence identity (ranging between 62%-73%), the nanobodies adopt distinct conformations (ranging between 3-7 Å all-atom RMSD) when bound to their targets. In the case of antibody-antigen series, the N-terminal domain of the SARS-CoV-2 nucleocapsid (N-) protein was bound to different human antibodies, derived from COVID-19-infected individuals. The antibodies’ light and heavy chains exhibited moderate to high sequence identity, ranging from 43% to 81% and 74% to 96%, respectively, leading to 4-6 Å all-atom RMSD variations. In addition to these cross-species and immunity complexes, the primary target sources were bacterial and eukaryotic cells. Five targets presented different conformational states that were induced by ligand binding, mutation, or buffer conditions. The heptameric (H1171) and octameric (H1172) Holliday junction targets showcase the ligand-binding-induced conformational changes happening in the corresponding hexameric apo structure T1170o. Target T1109o, an isocyanide hydratase mutant, is an example of conformational changes due to a one-point mutation (D183→A183) in the corresponding wild-type structure T1110o. The point mutation resulted in an all-atom RMSD change of 11.2 Å. The ancient protein reconstruction target T1160o exhibits conformational differences compared to what is available in the literature due to different compounds used in the crystallization buffer^62^. T1161o is a mutation-induced version of T1160o, designed to explore the connection between an ancient DPBB (DNA/RNA polymerase binding) protein and a RIFT (ribosomal) protein. Each monomer of this protein had five amino acid differences compared to the original sequence, leading to a 1.6 Å all-atom RMSD change.

**Table 1.**
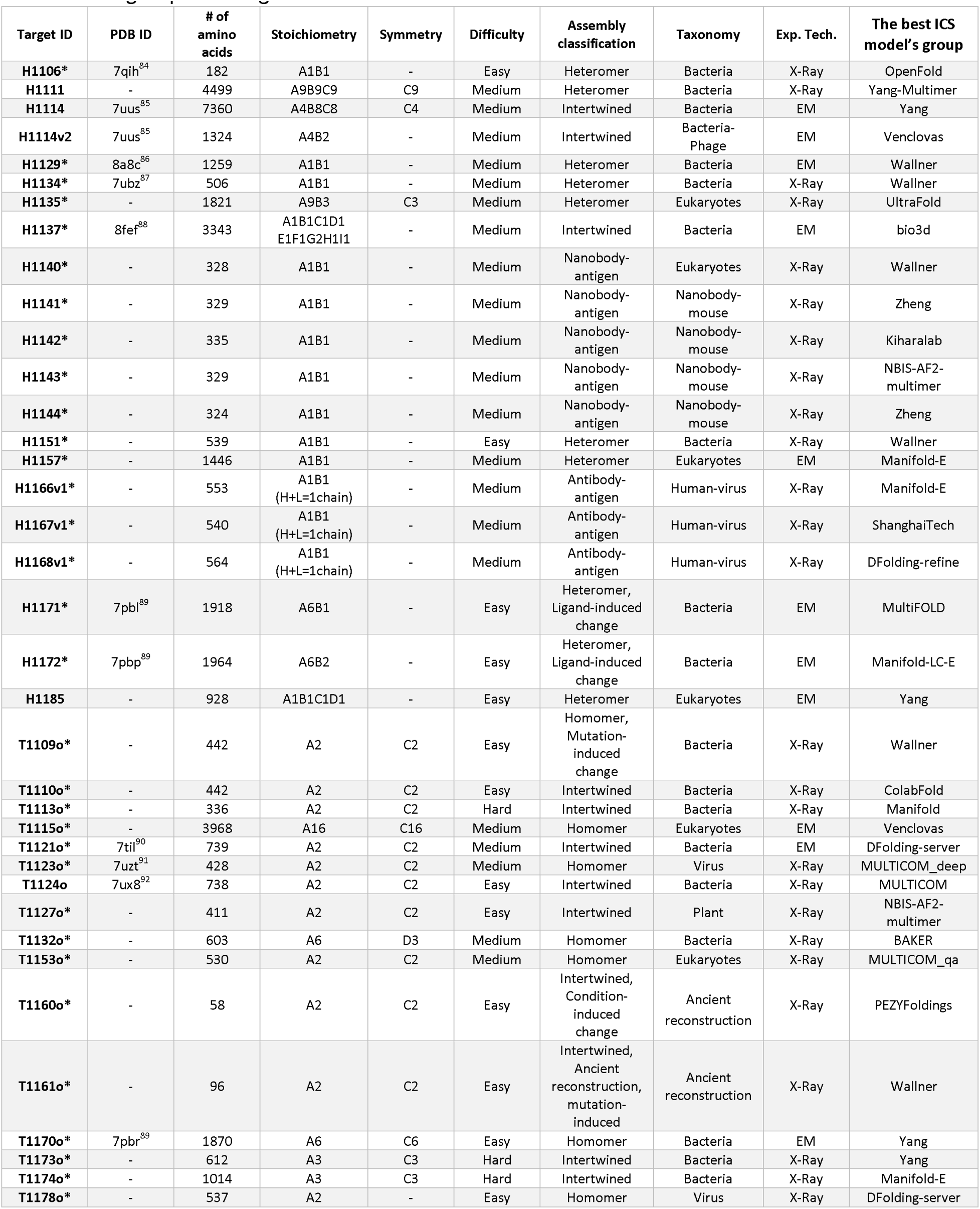
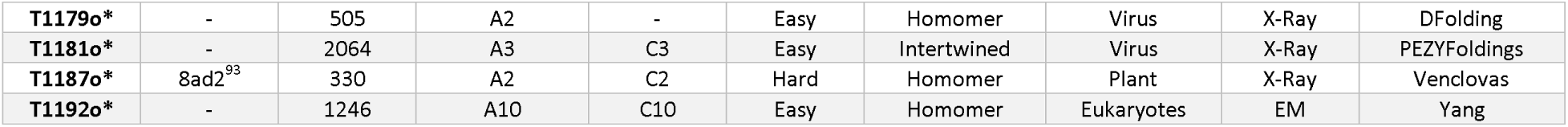
The CASP15 assembly targets with target’s pdb id (if available), size, stoichiometry, symmetry, prediction difficulty, assembly class, taxonomy, experimental technique used to resolve it, and the best ICS model’s group. The targets shared with CAPRI are indicated with *.

All targets were evaluated with all interfaces contributing to the assessment score. For one of the targets with complicated stoichiometry (H1114, A4B8C8), the prediction challenge was to predict the smallest interface of the complex, formed between an intertwined tube and a protein dimer. We, therefore, additionally evaluated this subcomplex (A4B2) as a separate target (H1114v2) (Figure S1B). This increased the total number of evaluated cases to 41. From the perspective of prediction difficulty, the targets were categorized into 16 easy, 21 medium, and 4 hard cases according to the principles outlined in Methods (see Table S1). Further details on the assembly targets are presented in Table 1, together with their structural depictions in Figure S2.

A record number of participants - 61 human and 26 server groups - took part in CASP15. These groups submitted a total of 11,000 models, presenting the load four times larger than that in the previous CASP (Table 2). All targets were shared with CAPRI, and most of them were selected for CAPRI prediction (Table 2). Among the 87 CASP15 participating groups, 60 submitted predictions on more than half of the targets. According to the CASP15 Abstract Book, 90% of the groups used the AF2M methodology in one way or another^37,47^. Some groups directly used the standard setup of AF2M, while others tried to enhance the sampling and/or scoring procedures of AF2M by supplying custom MSAs, templates, increasing the number of recycles used in sampling, or activating the dropout layers in the AI model to increase sampling diversification. Only very few groups performed solely classical docking and template-based modeling, but this time starting from the AF2 tertiary structure models. A small set of groups utilized normal mode analysis and short molecular dynamics to model conformational changes happening upon binding. A brief description of these methods can be found in the CASP15 Abstract Book^47^. Finally, it is worth mentioning that all AF2M-dependent CASP groups encountered a bottleneck in the straight-on modeling of large complexes, which had to be modeled upon dissecting the whole assembly into smaller subcomplexes.

**Table 2.** Statistics of CASP assembly rounds over the years, including the share of heteromeric targets, the number of targets having > 2 chains, difficulty class shares, number of targets shared with the joint CAPRI round, the ICS profiles acquired for the hard targets, the number of competitors and the total number of models submitted, the top performing approaches.

| CASP Round | # of targets (heteromers /all) | # of targets >2 chains | Difficulty (easy/ medium/ hard) | # of shared targets with CAPRI | Max. ICS observed for hard targets | # of groups/# of models | Notable approaches |
| --- | --- | --- | --- | --- | --- | --- | --- |
| CASP12 | 8/30 | 14 | 9/6/15 | 15 | 0.15 | 108/1600 | Homology modeling |
| CASP13 | 12/42 | 17 | 12/17/13 | 20 | 0.52 | 45/5000 | Homology modeling |
| CASP14 | 17/22 | 13 | 2/20/6 | 12 | 0.48 | 39/2500 | Homology modeling<br>Docking<br>DL-based contacts<br>Fold-and-dock |
| CASP15 | 20/40 | 17 | 14/22/4 | 36 | 0.95 | 87/11,000 | Mainly AF2-M based modeling |

### CASP15 witnessed an unprecedented improvement in assembly modeling

In CASP15, we employed four different metrics to evaluate the accuracy of assembly models (see Methods). Distributions of the evaluation scores for all targets are shown in Figure 1. As defined in Methods, we consider a model to be successful (or acceptable) if it scores above 0.50 according to all evaluation metrics. Strikingly, the CASP15 community could submit at least one successful model for 90% (37 out of 41) targets (Table 1). For comparison, only 31% of CASP14 targets had successful models. In CASP15, the failing targets correspond to two antibody-antigen complexes (H1166, H1167), one nanobody-antigen complex (H1142), and a viral homodimer (T1123o).

**Figure 1.**
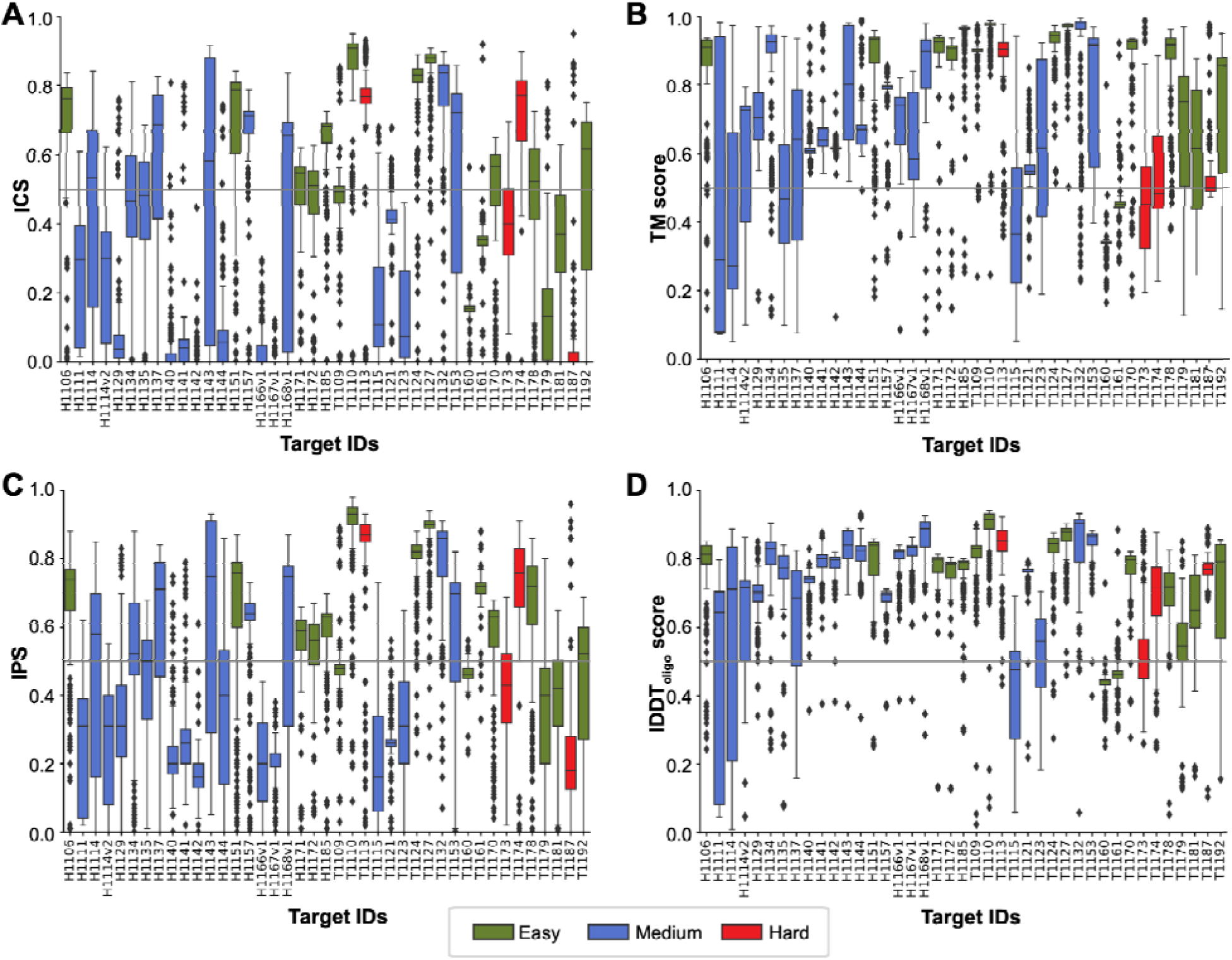
Per-target distribution of A. ICS, B. TM, C. IPS, and D. lDDT metrics calculated over the submitted models. The box coloring follows the target difficulty color code (green: easy, blue: medium, red: difficult). The threshold for a model to be successful (0.50) is marked with a horizontal gray line.

Best predictors managed to model the overall fold of CASP15 targets extremely well. According to the fold similarity measures (lDDT and TM), 100% of CASP15 targets had model scoring over 0.50 (76% in CASP14) and 90% scoring over 0.75 (14% in CASP14). Getting interface regions accurately proved to be a slightly more difficult task, with 90% of CASP15 targets having models scoring over 0.50 according to the interface-based metrics IPS and ICS (31% in CASP14). Figure 2A confirms the striking improvement in the accuracy of interface modeling in the latest CASP. While in the pre-CASP15 editions the vast majority of targets (around 70%, red) were falling in the poor accuracy zone and only very small fraction (around 10%, green) in the high accuracy zone, in CASP15 the situation flipped with only less than 10% of targets being modeled poorly and more than a half modeled with high accuracy. The mean ICS score of the top model doubled from 0.37 in CASP14 to 0.74 in CASP15, almost reaching the lower boundary of the high-accuracy prediction zone (0.75 - see Methods for zone definitions). Notably, best model for some of the hardest targets almost perfectly reproduced native interfaces in CASP15 (for example, ICS of 0.95 for T1187o or 0.90 for T1174o), while never crossing the 0.52 level in all previous CASP rounds (Table 2).

**Figure 2.**
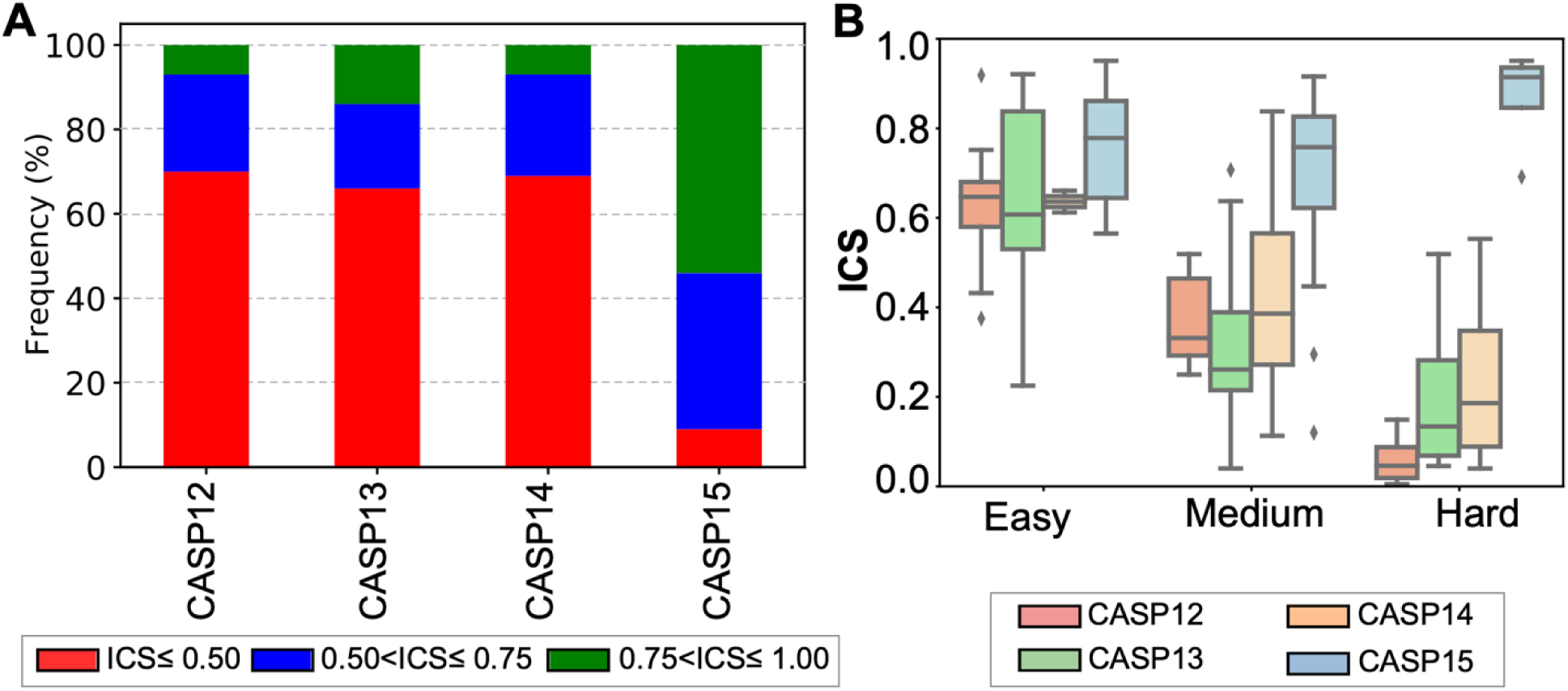
**A. Percentage of the best models in CASP12-15 falling into three interface accuracy bins:** poor: ICS<0.50, red; good: 0.50 ≤ ICS < 0.75, blue; and high: ICS ≥ 0.75, green. **B. Box-and-whisker plots for the best ICS scores generated in each round.** The distributions are presented under different difficulty categories.

In CASP15, half of our targets were homomers. Also, the majority of the targets had complete or partial templates, de-facto making these targets easier for modeling. To ensure that the advancement observed in CASP15 was not solely due to the decreased difficulty of the target set, we conducted a difficulty-tuned comparative analysis of results across different CASP rounds (Figure 2B, Methods). This analysis demonstrated a significant performance enhancement across all difficulty categories, with an upward shift of 0.2 ICS for the easiest targets and 0.5 for the hardest ones, which shows that target difficulty was not the major determinant in the unprecedented accuracy improvement in CASP15.

### Best performing CASP15 groups and their modeling strategies

We ranked participating groups according to a Z-score-based function composed of equally weighted inputs from ICS, IPS, lDDT, and TM metrics (Eq.5, Methods, Table S2). Zheng group ranked at the top (Figure 3A), followed by the second tier of groups including Venclovas and Wallner groups, and the third tier comprising Yang and Yang-Multimer. The robustness of the ranking is confirmed by the jackknife cross-validation. If we rank groups on the best model submitted on targets, the first two tiers would remain the same, while the third tier would include PEZYFolding and Kiharalab. Zheng, Venclovas, and Kiharalab groups predicted all 41 targets, Wallner - 36 targets, and PEZYFolding and Yang - 39. If we consider the average Z-scores for ranking, Zheng continues to lead as the top-performing group, followed by Wallner and Venclovas (Figure S3). These groups achieved successful model submissions (considering their best ICS models) for a substantial percentage of the targets they participated in: 73% for Zheng and Venclovas, 69% for Wallner, 64% for Yang, 67% for PEZYFolding, and 56% for KiharaLab. The NBIS-AF2M group, which employed the standard AF2M settings (Methods), ranked 30^th^ and successfully generated at least one good ICS model for 54% of the targets (22/41).

**Figure 3.**
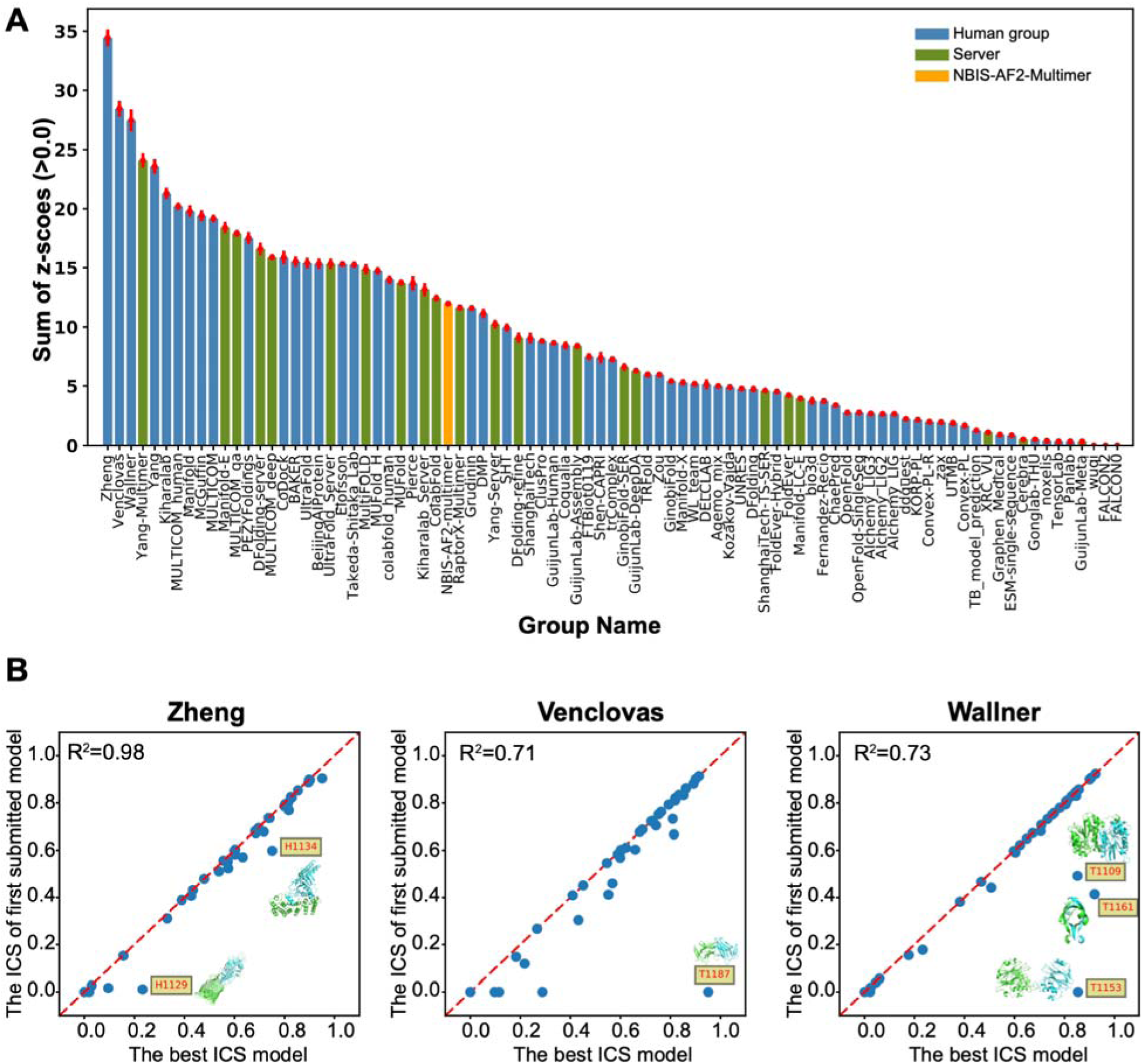
A. The CASP15 assembly groups, ordered from left to right according to their relative performance. For the final ranking, we utilize the average scores obtained from the jackknifing protocol (Methods). The bar height represents the cumulative Z-score (Eq.7), while the error whiskers mark the standard deviation of the jackknifed data set. Human groups are shown in blue, servers in green, and the baseline NBIS-AF2-Multimer in orange. **B. Model ranking efficiency of the top performing groups.** The ICS of the first submitted model is plotted against the best ICS model. Yellow boxes highlight off-diagonal targets (i.e., those, for which predictors wrongly identified their best model).

The top-ranked Zheng group utilized DeepMSA2 pipeline for generating MSAs that are subsequently fed into AF2M^63^. The MSAs were built by iteratively running HHblits^64^, Jackhmmer^65^, and HMMsearch^65^ on several databases. While modeling heteromers, they sampled all possible MSA pairing combinations to obtain the most optimal pairing scenario. Models were ranked according to the AF2M’s pTM score. The Venclovas group used ColabFold^32^, which employs MMseqs2 to search genetic databases and construct MSA^66^. For sampling they used both the AF2M and monomeric AF2 AI models (so, both versions 2.1^26^ and 2.2^33^). They sampled different paired MSAs by using different sequence databases. They also increased the number of recycles. When ColabFold failed to assemble the complex (due to the system size limitations), Venclovas’ team used rigid body docking. SAM was used to dock symmetric homomers, while FTDOCK and HEX were used to assemble heteromers^67–69^. After pooling all models, they ranked the models with their consensus VoroIF-jury method^70,71^. Wallner sampled an exhaustive set of models by using AF2M versions 2.1^26^ and 2.2^33^. While using these AI models, they activated the dropout layers in the Evoformer block, increased the number of recycles, as well as the number of seeds, and turned on/off the template usage. They ranked the sampled conformations according to the AF2M’s model confidence score (Methods). Wallner aimed to submit a diverse set of models having a high model confidence score (>0.7)^72^. The Yang teams replaced AF2M’s template search with HHsearch and disabled MSA pairing^73^. For model ranking, they used the same scoring scheme as Wallner. PEZYFolding and Kiharalab groups performed well on sampling, but not on scoring. PEZYFolding used custom MSAs as an input to AF2M. For this, they used their sequence search tools PZLAST and PSI-BLASTexB next to AF2M’s sequence search tools^74–76^. These tools were run over several public and private sequence databases. The top model was picked either by plDDT or ipTM. The rest of the models were selected to reflect minimum structural similarity among all submitted models. Kiharalab generated its custom MSA by running HHblits, Jackhmmer over different genetic databases in an iterative manner. For the symmetric homomeric complexes, where AF2M failed, they used the docking tool SAM, like the Venclovas group. SAM was run with the bound monomeric conformations obtained from AF2M.

To examine the sampling space covered by the top five assembly groups, we analyzed the similarity of their ICS scores compared to the baseline AF2M’s (Figure S4). Zheng and Yang-Multimer showed the highest correlations with AF2M, with values of 0.78 and 0.74, respectively. On the other hand, Wallner and Venclovas had the least similar models, with correlations of 0.43 and 0.51, respectively. This difference could be attributed to Wallner and Venclovas focusing on diversifying their final model sets. To evaluate the effectiveness of these groups’ ranking schemes, we compared their best ICS models with their top-ranking (i.e., first) model’s ICS (Figure 3B). According to our analysis, the pTM scoring method worked exceptionally well for Zheng. Wallner performed as the second-best scorer, although they couldn’t rank their best models for three cases (T1109o, T1161o, T1153o). Lastly, Venclovas had only one significant discrepancy between what they thought was their best model and their actual best ICS model (T1187o). Strikingly, for the other targets, their custom scoring function performed almost as well as AF2M’s confidence score.

Homology modeling and docking driven by AF2 tertiary structure models was employed as the main methodology in CASP15 only by a few groups (ChaePred, ClusPro, and FTBiot0119). These groups achieved lower rankings than standard AF2M, indicating that simultaneous modeling of monomers in the context of an assembly is a more efficient modeling practice than these semi-classical approaches.

### Exploring the factors affecting model accuracy: MSA, templates, and sampling parameters

In this section we investigate the influence of input data and sampling parameters on the accuracy of protein complexes generated by the AF2M AI model. First, we analyze whether richness of evolutionary information (in terms of the complex’ combined MSA depth) has an impact on the accuracy of assembly models. Thirty out of 41 assembly targets are suitable for this analysis; the remaining 11 are disqualified as either representing inter-species complexes or being merely a duplication of other targets. Complexes H1129, H1166-H1168, and H1140-H1144 are cross-species by their nature and as such do not contain intra-species binding signal in their individual subunit MSAs (i.e., for these targets the model accuracy will not be a function of their multimeric Neff). Targets T1109o and T1161o are close mutants of their wild-type relatives (T1110o and T1160o) that are already included in the selected 30 and contribute to the MSA-centered analysis. These eleven special interaction cases are examined further below from the perspectives of template availability and optimized sampling.

Figure 4A shows the relationship between the multimeric MSA depth, Neff (see Methods), and the best ICS scores for the qualified 30 targets, after their separation into three difficulty classes. The results reveal that the easiest for modeling targets have the deepest MSA (average Neff of 4203), while the most difficult ones have the shallowest MSA (average Neff of 1344). Interestingly, and against the expectation, interfaces of the hard modeling targets were modeled most accurately (average ICS of 0.87) in the hardest targets. A possible reason for this can be that all hard targets are rather small and formed by obligate interactions (Figure S5). The trend for a better performance with increasing alignment depth is apparent when analyzing the ICS-Neff correlation inside each target difficulty bin. These results are in line with the conclusions drawn for the tertiary structure prediction category^26,77,78^. Figure 4A also suggests that size of the complex may be another factor influencing modeling accuracy, as larger targets (1800 residues or more) were modeled slightly worse than the smaller ones (average ICS of 0.68 versus 0.75 for the latter).

**Figure 4.**
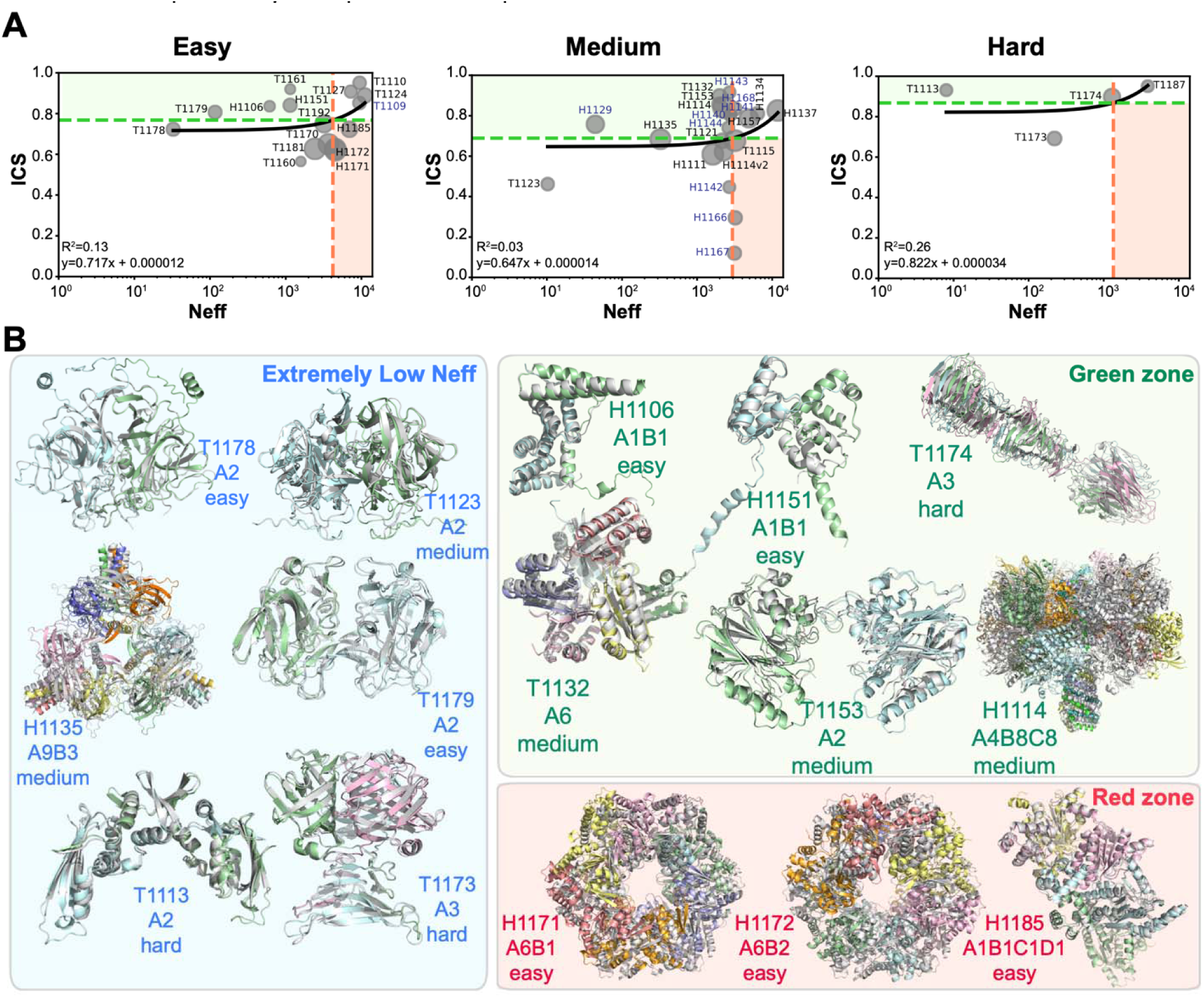
A. Relationship between baseline Neff values and the best ICS values for each difficult category. In all categories, the horizontal green dotted line represents the mean best-ICS value, and the vertical red dotted line represents the mean Neff value. The green zone represents areas where lower Neff values are associated with higher ICS values, while the red zone represents areas where higher Neff values are associated with lower ICS values. Neff values are depicted on the x-axis using a logarithmi scale to emphasize distinctions among low Neff values. The special cases (11 targets) are shown in purple. Markers of data points are scaled to their assembly size with larger complexes shown as larger circles. **B. Comparison of the best ICS models with the experimental structures.** The best ICS models (shown in color) superimposed on target structures (gray) for the green, red, and extremely low Neff zone targets (see panel A). Stoichiometry and difficulty definitions are provided for each target.

To examine the relationship between the model accuracy and Neff in more detail, we divided the plot in Figure 4A into four distinct zones based on the mean ICS and Neff values. The mean ICS are 0.77, 0.69 and 0.87 and the mean Neff are 4203, 2952 and 1344 for easy, medium, and hard classifications, respectively. We focused on two zones: the “green” zone, where models with higher-than-average ICS scores could be generated despite relatively low Neff values, and the “red” zone, where lower ICS models were generated despite relatively high Neff values. The green zone includes three targets in the easy category (H1106, H1151, and T1179 -all small dimers), three bacterial symmetric cases in the medium category (T1132, T1153, and H1114), and two intertwined homomers in the hard category (T1113o and T1174o) (Figure 4B). There is a tendency for the green zone targets to be smaller or composed of intertwined molecules. The red zone is populated only for easy targets, with all cases being large-sized targets (H1172, H1171, and H1185). We also paid special attention to the targets with good or high accuracy ICS models and extremely low Neff values (i.e., points in immediate closeness to the Y-axis). These cases include T1178o, T1179o (small viral homodimers with structurally distant templates), H1135 (a cyclic homononamer in complex with three repeating peptides having a poor template), and T1113o, T1173o (intertwined bacterial homomers with no templates at the sequence and structure level) (Figure 4B). As these complexes are all made of obligate interactions, AF2M might have tried to assemble them by using building blocks learned from various entries in the Protein Data Bank, even in the absence of the MSA information. This observation is also supported by the fact that intertwined complexes could be predicted accurately even in the absence of MSA pairing, as proved by Yang. The absence of homologous sequences had a substantial negative impact only in the case of T1123o, where the interface was primarily composed of loop interactions.

Among the 11 special case targets, two complexes, H1143 and H1168, had templates highlighting the antigen binding site on the target. H1143’s template was the substrate bound CNPase (Figure S6A, Table S1). H1168v1’s template was another antibody-binding N-protein (Figure S6B, Table S1). For both targets, the availability of templates resulted in the generation of numerous high-accuracy models (Figure 1A). The remaining complexes had partial or complete templates only at the monomeric level. Among these, exhaustive sampling was necessary to produce differentially improved models for the mutation-induced T1109o and T1161o, the host-pathogen H1129, and nanobodies H1140-H1141-H1143, compared to the rest of the community (see positive outliers generated for these complexes in Figure 1A). Among the top-performing groups, Wallner and Venclovas stood out for their utilization of exhaustive sampling techniques. The Wallner group notably produced the best or one of the best high accuracy models for targets T1109o, T1161o, H1129, H1140, H1141, and H1144 (Figure S6). The Venclovas group produced the second-best ICS model for target H1129. Interestingly, in the case of two nanobody-antigen targets (H1141 and H1144), the custom MSA approach by the Zheng group worked and yielded the best ICS models. Only H1142 remained elusive for prediction by all groups, with the best model achieving an ICS score of 0.45 (Figure S6A). The buried surface area for these nanobody-antigen complexes follows the order: 1174.1 Å2 (H1142) < 1545.3 Å2 (H1140) < 1784.1 Å2 (H1144) < 1854.1 Å2 (H1141). So, the smallest interface was observed in H1142, hinting at a transient interaction, and thus posing a challenge for its prediction. Nevertheless, predicting real antibody-antigen interactions in the absence of templates (H1166-H1167) remains a challenge (Figure S6B).

### Advancing Beyond Standard AF2M Modeling

The community submitted at least one acceptable model for 90% of the targets, while the baseline predictor (NBIS-AF2-M) managed to generate an acceptable model only in 54% of cases. The performance gap observed between NBIS-AF2-M and the community motivated us to conduct further analysis to determine the factors that contributed to the improvements achieved by top ranking groups compared to the standard AF2M pipeline. For this, we identified the best ICS model for each target and calculated the difference between its ICS score and that of the best NBIS-AF2-M model. Figure 5A shows that the community significantly outperformed AF2M (ΔICS>0.25) on 15 targets (T1109o, T1115o, T1121o, T1160o, T1161o, T1179o, T1187o, H1111, H1114, H1114v2, H1129, H1135, H1140, H1141, H1144). Among these, enhanced sampling might explain the improvement for T1109o, T1161o, H1129, H1140, H1141, H1144, as discussed above. H1111, H1114, H1114v2, T1115o, H1135 share a common characteristic — they are large targets containing more than 1800 amino acids (Table 1, Figure S2). Modeling the complete stoichiometry of these complexes was beyond the capacity of the AF2M server and therefore for these complexes, AF2M submitted a subcomplex of the entire assembly. As an example, H1111 represents a symmetric A9B9C9 bacterial assembly. The best model for this target was submitted by the Yang group, achieving an ICS of 0.61 and a ΔICS of 0.57. NBIS-AF2-M, in contrast, submitted a trimeric model where the largest subunit hindered the formation of the 27-meric ring (Figure 5B). As another example, the bacterial 20-mer assembly H1114 features an intertwined tube at its center, which could not be built in the absence of partners (Figure 5C). Yang submitted the best model for the entire complex with an ICS of 0.84 and a ΔICS of 0.67. Two more targets are worth further exploring. The first one is T1179o, where AF2M predicted a more compact model than the actual structure. For this case, an unpaired MSA-driven approach, DFolding-server produced a model with an ICS of 0.81 and a ΔICS of 0.67 (Figure 5D). The other one is H1129, where the complex was left undocked by the AF2M but was accurately predicted by Wallner as described above (Figure 5E). Finally, we also observed that the community and baseline AF2M methods performed similarly on all the Neff-based green and red zone complexes, except for H1114 (Figure 4A). For the template driven H1143 and H1168 complexes too, the community could not produce better models than the AF2M (Figure S6B).

**Figure 5.**
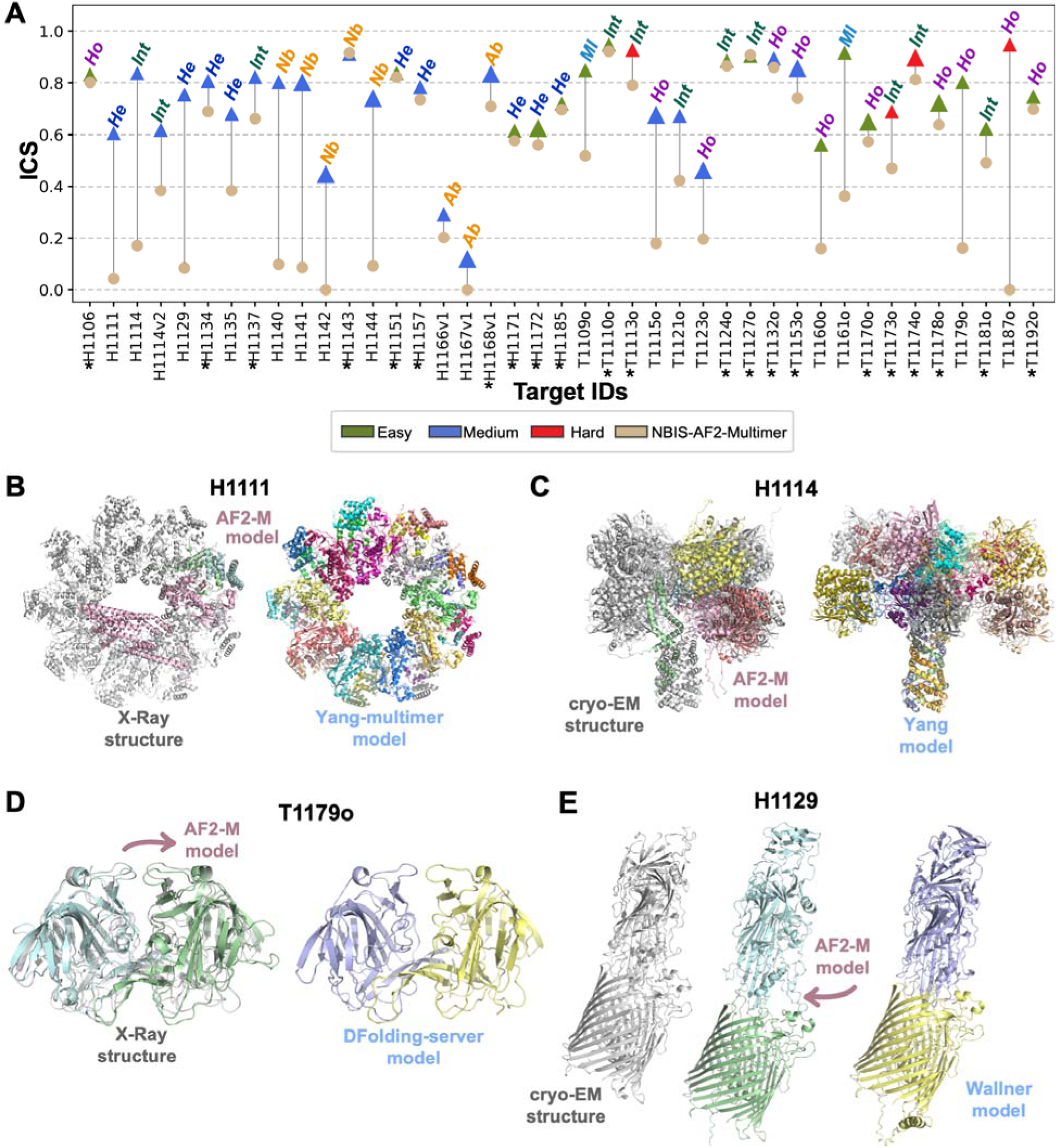
A. The best ICS model submitted by the community vs. the best ICS model submitted by NBIS-AF2-M. The color code used for the best community submissions follows the one in Figure 1. The best NBIS-AF2-M models are shown in tan. Each data point is marked with the complex type, where Ho stands for homomer, He for heteromeric, Int for intertwined, Nb for nanobody-antigen, Ab for antibody-antigen, and MI for mutation-induced change. The targets where NBIS-AF2-M and the community produced similar successful models (ΔICS<0.25) are marked with an asterisk *. The tabulated form of this plot is provided as Table 3. **B. The best ICS model submitted by the community, the best ICS model submitted by AF2M and the experimental structure for H1111, C. H1114, D. T1179o and E. H1129.** The best ICS models were depicted in color and the experimental structures were depicted in gray.

**Table 3.** ΔICS values for each target, the best ICS of NBIS-AF2-M, the best ICS and its performed group, which groups performed well and the reason for NBIS-AF2-M failure.

| Target ID | NBIS-AF2-M ICS | The best ICS | The best ICS model's group | ΔICS | Community and NBIS-AF2-M performed well | Nobody performed well | Community performed well | The reason of NBIS-AF2-M failure |
| --- | --- | --- | --- | --- | --- | --- | --- | --- |
| H1106 | 0.80 | 0.84 | OpenFold | 0.03 | x |  |  |  |
| H1111 | 0.04 | 0.61 | Yang-Multimer | 0.57 |  |  | x | technical limitation |
| H1114 | 0.17 | 0.84 | Yang | 0.67 |  |  | x | technical limitation |
| H1114v2 | 0.38 | 0.62 | Venclovas | 0.24 |  |  | x | technical limitation |
| H1129 | 0.08 | 0.76 | Wallner | 0.67 |  |  | x | wrong interface |
| H1134 | 0.69 | 0.81 | Wallner | 0.12 | x |  |  |  |
| H1135 | 0.38 | 0.68 | UltraFold | 0.30 |  |  | x | wrong interface |
| H1137 | 0.66 | 0.83 | bio3d | 0.16 | x |  |  |  |
| H1140 | 0.10 | 0.81 | Wallner | 0.71 |  |  | x | Nanobody-antigen |
| H1141 | 0.09 | 0.80 | Zheng | 0.72 |  |  | x | nanobody-antigen |
| H1142 | 0.00 | 0.45 | Kiharalab | 0.45 |  | x |  | nanobody-antigen |
| H1143 | 0.92 | 0.92 | NBIS-AF2-multimer | 0.00 | x |  |  |  |
| H1144 | 0.09 | 0.74 | Zheng | 0.65 |  |  | x | nanobody-antigen |
| H1151 | 0.82 | 0.84 | Wallner | 0.02 | x |  |  |  |
| H1157 | 0.74 | 0.79 | Manifold-E | 0.05 | x |  |  |  |
| H1166v1 | 0.20 | 0.30 | Manifold-E | 0.09 |  | x |  | antibody-antigen |
| H1167v1 | 0.00 | 0.12 | ShanghaiTech | 0.12 |  | x |  | antibody-antigen |
| H1168v1 | 0.71 | 0.84 | DFolding-refine | 0.13 | x |  |  |  |
| H1171 | 0.58 | 0.62 | MultiFOLD | 0.04 | x |  |  |  |
| H1172 | 0.56 | 0.63 | Manifold-LC-E | 0.07 | x |  |  |  |
| H1185 | 0.70 | 0.72 | Yang | 0.03 | x |  |  |  |
| **T1109o** | 0.52 | 0.85 | Wallner | 0.33 |  |  | x | Mutation induced change |
| **T1110o** | 0.92 | 0.95 | ColabFold | 0.03 | x |  |  |  |
| **T1113o** | 0.79 | 0.93 | Manifold | 0.14 | x |  |  |  |
| **T1115o** | 0.18 | 0.68 | Venclovas | 0.50 |  |  | x | technical limitation |
| **T1121o** | 0.42 | 0.68 | DFolding-server | 0.25 |  |  | x | wrong interface |
| **T1123o** | 0.20 | 0.46 | MULTICOM_deep | 0.27 |  | x |  | viral protein/weak evolutionary signal |
| **T1124o** | 0.87 | 0.89 | MULTICOM | 0.02 | x |  |  |  |
| **T1127o** | 0.91 | 0.91 | NBIS-AF2-multimer | 0.00 | x |  |  |  |
| **T1132o** | 0.86 | 0.90 | BAKER | 0.04 | x |  |  |  |
| **T1153o** | 0.74 | 0.86 | MULTICOM_qa | 0.12 | x |  |  |  |
| **T1160o** | 0.16 | 0.57 | PEZYFoldings | 0.41 |  |  | x | Mutation induced change |
| **T1161o** | 0.36 | 0.92 | Wallner | 0.56 |  |  | x | Mutation induced change |
| **T1170o** | 0.57 | 0.65 | Yang | 0.08 | x |  |  |  |
| **T1173o** | 0.47 | 0.69 | Yang | 0.22 | x |  |  |  |
| **T1174o** | 0.81 | 0.90 | Manifold-E | 0.09 | x |  |  |  |
| **T1178o** | 0.64 | 0.72 | DFolding-server | 0.09 | x |  |  |  |
| **T1179o** | 0.16 | 0.81 | DFolding | 0.65 |  |  | x | wrong interface |
| **T1181o** | 0.49 | 0.63 | PEZYFoldings | 0.14 | x |  |  |  |
| **T1187o** | 0.00 | 0.95 | Venclovas | 0.95 |  |  | x | wrong interface |
| **T1192o** | 0.70 | 0.75 | Yang | 0.05 | x |  |  |  |

### From inter-monomer to interdomain predictions

In CASP15, the assessment of interdomain interface predictions was conducted alongside the assembly category, as both categories share conceptual similarities and can be analyzed using similar assessment approaches.

Tertiary structure prediction in CASP was evaluated at the level of evaluation units (EU), which are either structural domains or easily predictable combinations of such (see Kryshtafovych and Rigden for details of the CASP15 EUs definition)^79^. The tertiary structure assessment concentrates on the question of how well structures of separate domains are reproduced, while the interplay between domains is left outside the scope of that assessment and is the subject of this study. All in all, 22 single-sequence CASP15 targets (monomers and subunits of multimers) were split into multiple EUs (called here for simplicity “domains”). We took these targets, calculated their interdomain interactions, and kept the domain pairs having at least ten interdomain contacts (see Methods). In the end, we evaluated 17 interfaces from 7 targets (Table 5). To assess accuracy in modeling interdomain interfaces, we employed the ICS, IPS, and QS_global_ metrics. Similar to the assembly category, a successful model was defined as having scores above 0.50 for all metrics. According to this definition, 12 out of the 17 interfaces had successful predictions. However, five interfaces (T1125-D12, T1125-D34, T1125-D56, T1165-D16, and T1169-D12) proved to be challenging for the community (Figure 6). For one of these interfaces, T1125-D56, two of the evaluation scores were well above 0.50 (the best ICS of 0.63, and the best IPS of 0.67), while the third (QS_globa_l) was slightly below 0.50. We decided to count this target also as a successful case, thus bringing the total number of successful predictions to 13 and achieving an overall success rate of 76.5%.

**Figure 6.**
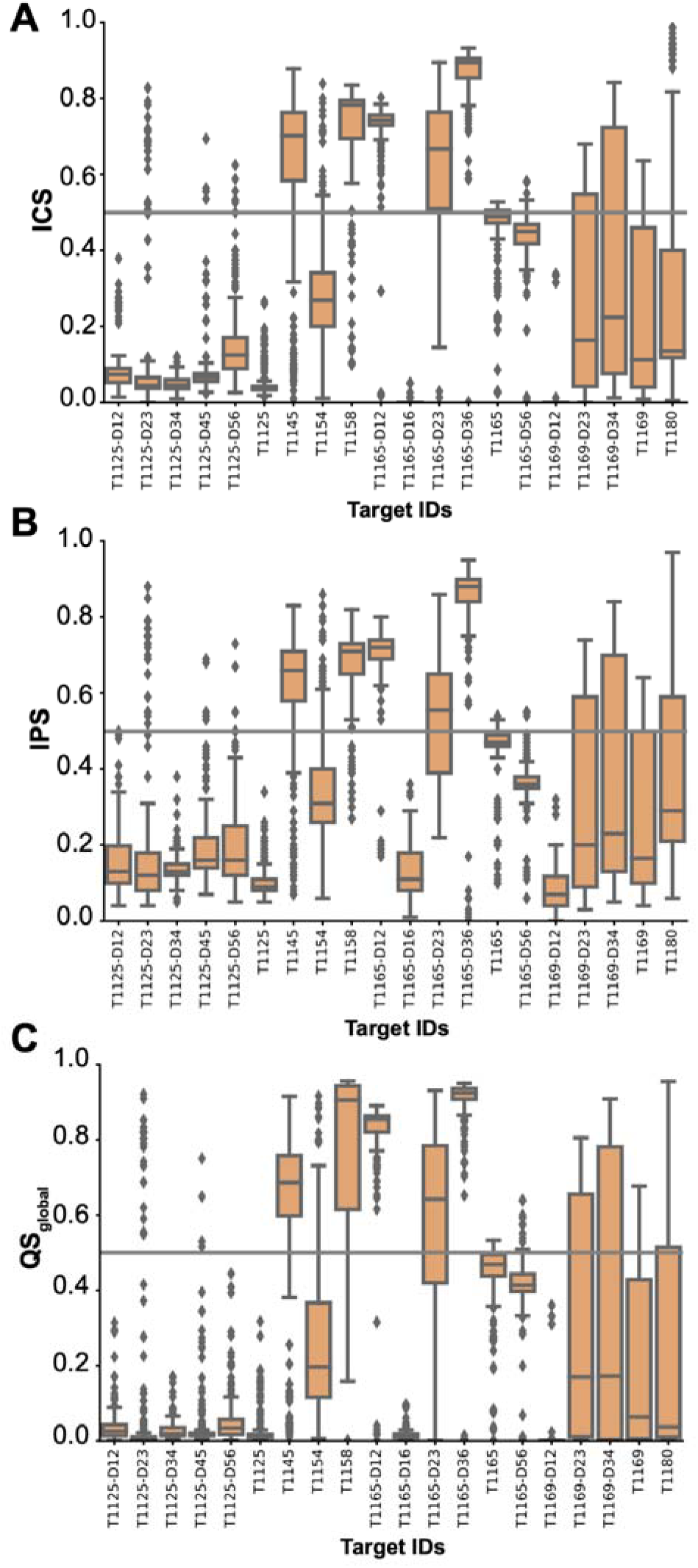
Per-target distributions of A. ICS, B. IPS and C. QS_global_ scores on interdomain targets. The 0.50 solid gray line mark denotes the threshold for a successful prediction.

**Table 4.** Assembly Neff-multimer and subunit Neff values calculated for the MSA obtained by standard AF2-M sampling. s denotes subunit and D denotes domains. Neff-multimer calculation is performed as described in Methods.

| Assembly target ID | Subunit target ID | Neff of subunit | Neff-multimer |
| --- | --- | --- | --- |
| H1106 |  |  | 612 |
|  | T1106s1 | 246 |  |
|  | T1106s2 | 1522 |  |
| H1111 |  |  | 1617 |
|  | T1111s1 | 245 |  |
|  | T1111s2 | 1490 |  |
|  | T1111s3 | 11563 |  |
| H1114 |  |  | 2250 |
|  | T1114s1-D1 | 241 |  |
|  | T1114s2 | 5478 |  |
|  | T1114s3 | 8626 |  |
| H1114v2 |  |  | 2250 |
|  | T1114s1-D1 | 241 |  |
|  | T1114s2 | 5478 |  |
|  | T1114s3 | 8626 |  |
| H1129 |  |  | 44 |
|  | T1129s2 | 44 |  |
| H1134 |  |  | 6372 |
|  | T1134s1 | 9879 |  |
|  | T1134s2 | 4110 |  |
| H1135 |  |  | 324 |
|  | T1135s1 | 3875 |  |
|  | T1135s2 | 27 |  |
| H1137 |  |  | 11889 |
|  | T1137s8 | 12171 |  |
|  | T1137s9 | 11613 |  |
| H1140 |  |  |  |
|  | H1140s1 | 672 | 2778 |
|  | H1140s2 | 11493 |  |
| H1141 |  |  | 2754 |
|  | H1141s1 | 672 |  |
|  | H1141s2 | 11290 |  |
| H1142 |  |  | 2682 |
|  | H1142s1 | 672 |  |
|  | H1142s2 | 10714 |  |
| H1143 |  |  | 2711 |
|  | H1143s1 | 672 |  |
|  | H1143s2 | 10940 |  |
| H1144 |  |  | 2597 |
|  | H1144s1 | 672 |  |
|  | H1144s2 | 10046 |  |
| H1151 |  |  | 1142 |
|  | T1151s2 | 1142 |  |
| H1157 |  |  | 4889 |
|  | T1157s1-D1 | 6142 |  |
|  | T1157s2-D1 | 5225 |  |
|  | T1157s2-D2 | 2853 |  |
|  | T1157s2-D3 | 6239 |  |
| H1166v1 |  |  | 3215 |
|  | H1166s1 | 14760 |  |
|  | H1166s2 | 12892 |  |
|  | H1166s3 | 175 |  |
| H1167v1 |  |  | 3138 |
|  | H1167s1 | 14571 |  |
|  | H1167s2 | 12142 |  |
|  | H1167s3 | 175 |  |
| H1168v1 |  |  | 3166 |
|  | H1168s1 | 14246 |  |
|  | H1168s2 | 12756 |  |
|  | H1168s3 | 175 |  |
| H1171 |  |  | 4521 |
|  | T1171s1 | 12756 |  |
|  | T1171s2 | 1602 |  |
| H1172 |  |  | 4521 |
|  | T1171s1 | 12756 |  |
|  | T1171s2 | 1602 |  |
| H1185 |  |  | 6963 |
|  | T1185s1 | 9210 |  |
|  | T1185s2 | 8754 |  |
|  | T1185s4 | 4187 |  |
| T1109o |  |  | 9426 |
|  | T1109-D1 | 9426 |  |
| T1110o |  |  | 9478 |
|  | T1110 | 9478 |  |
| T1113o |  |  | 8 |
|  | T1113 | 8 |  |
| T1115o |  |  | 3201 |
|  | T1115-D1 | 4502 |  |
|  | T1115-D2 | 2276 |  |
| T1121o |  |  | 2160 |
|  | T1121-D1 | 1535 |  |
|  | T1121-D2 | 3040 |  |
| T1123o |  |  | 10 |
|  | T1123-D1 | 10 |  |
| T1124o |  |  | 11132 |
|  | T1124 | 11132 |  |
| T1127o |  |  | 7245 |
|  | T1127 | 7245 |  |
|  | T1127 | 7245 |  |
| T1132o |  |  | 1992 |
|  | T1132 | 1992 |  |
| T1153o |  |  | 1950 |
|  | T1153-D1 | 1950 |  |
| T1160o |  |  | 1584 |
|  | T1160 | 1584 |  |
| T1161o |  |  | 1148 |
|  | T1161 | 1148 |  |
| T1170o |  |  | 3686 |
|  | T1170-D1 | 3810 |  |
|  | T1170-D2 | 3567 |  |
| T1173o |  |  | 221 |
|  | T1173-D1 | 2870 |  |
|  | T1173-D2 | 17 |  |
| T1174o |  |  | 1288 |
|  | T1174-D1 | 3687 |  |
|  | T1174-D2 | 450 |  |
| T1178o |  |  | 33 |
|  | T1178 | 33 |  |
| **T1179o** |  |  | 117 |
|  | **T1179** | 117 |  |
| **T1181o** |  |  | 2432 |
|  | **T1181-D1** | 1191 |  |
|  | **T1181-D2** | 4965 |  |
| **T1187o** |  |  | 3859 |
|  | **T1187** | 3859 |  |
| **T1192o** |  |  | 3206 |
|  | **T1192s1** | 3206 |  |

**Table 5.** The CASP15 interdomain target list with target’s pdb id (if available), taxonomy, the experimental technique used to resolve its structure, number of domains, evaluated interfaces* and Buried Surface Area. *For domains, refer to Figure S7.

| Target ID | Pdb ID | Taxonomy | Exp. Tech. | Number of domains | Evaluated interfaces* | BSA (Å<sup>2</sup>) |
| --- | --- | --- | --- | --- | --- | --- |
| **T1125** | - | virus | X-Ray | 6 | D12, D23, D34, D45, D56 | 871, 894, 854, 937, 187 |
| **T1145** | - | bacteria | X-Ray | 2 | D12 | 516 |
| **T1154** |  | archaea | EM | 2 | D12 | 441 |
| **T1158** |  | eukaryotes | EM | 2 | D12 | 2231, 3072, 2927, 2799, 4932 |
| **T1165** |  | eukaryotes | EM | 6 | D12, D16, D23, D36, D56 | 1245, 894, 735, 1088, 815 |
| **T1169** | 8fjp<sup>94</sup> | eukaryotes | EM | 4 | D12, D23, D34 | 1685, 2381, 2115 |
| **T1180** | - | bacteria | X-Ray | 2 | D12 | 639 |

Predictors in the interdomain category were ranked according to the Z-score-based function presented in (Eq.9). UM-TBM and Yang-server groups outperformed other groups by a large margin (Figure 7A and S8, Table S3). The UM-TBM, developed by Zheng’s group, is a fully automated pipeline designed to generate tertiary structure models. This approach combine multiple tools and strategies, including contemporary deep learning approaches to predict contacts, angles, distances, as well as classical techniques, such as template-based modeling and molecular dynamics simulations. In particular, UM-TBM produced exceptionally good results for T1125-D23 (Table S4, Figure 7B). The second-best predictor, Yang-server, employed trRosettaX2 and AF2. Their pipeline incorporated the attention-based network from AF2 to improve the prediction of inter-residue distances and orientations, along with energy minimization from trRosetta. Notably, the Yang-server achieved top rankings in T1169 and T1154 targets (Table S4, Figure 7C, D).

**Figure 7.**
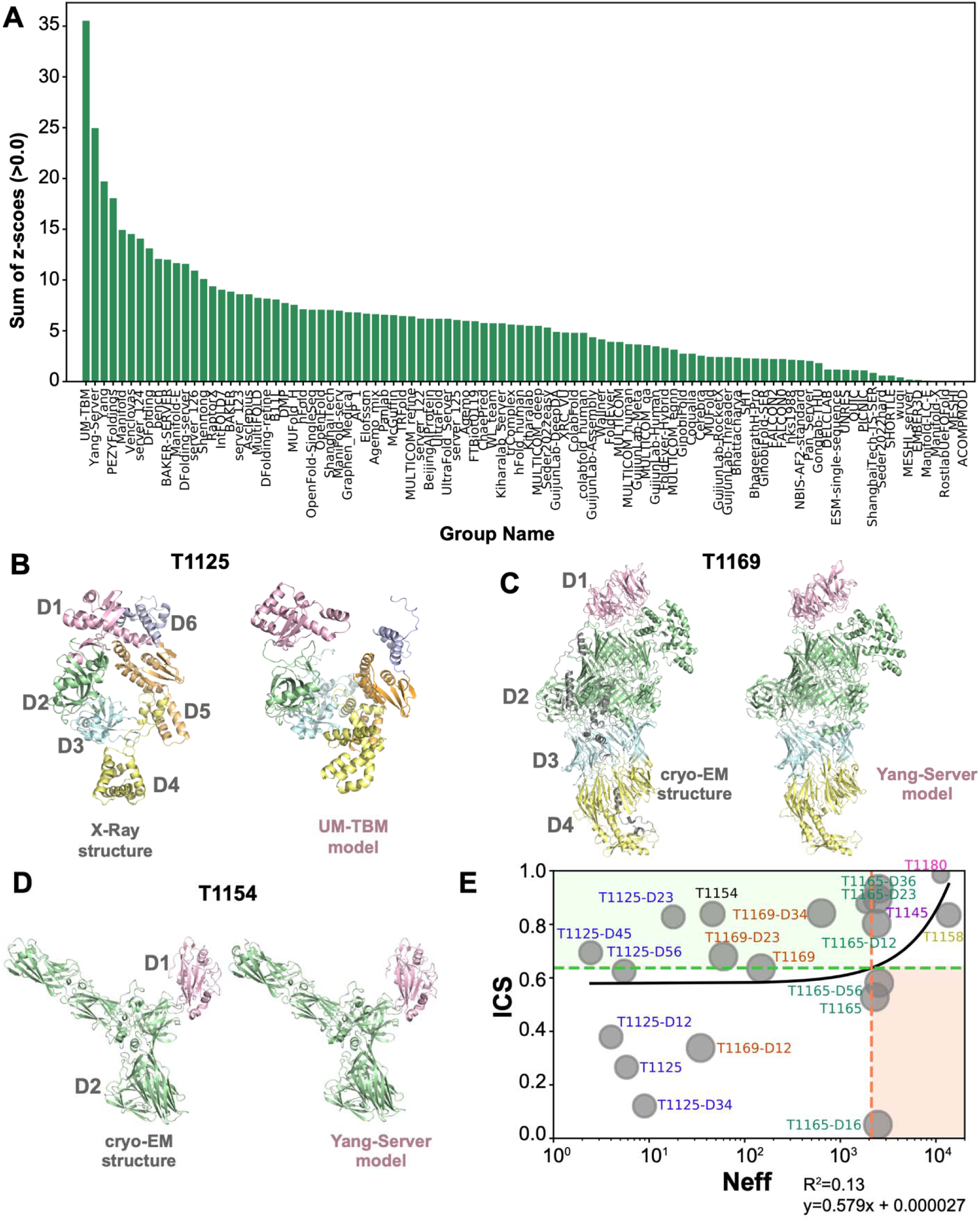
A. Performance of CASP15 groups in predicting interfaces of multi-domain targets. **The groups** are ranked according to their first submitted models with the Z-score function given in Eq.9. **B. The domain organization of T1125.** The top-ranking model generated for this target is submitted by UM-TBM. In this model, D6 is not closing upon D1, as in the crystal structure. **C. The domain organization of T1165.** The top-ranking model generated for this target is submitted by Yang-server. In this model, D1 is not closing upon D2, as in the cryo-EM structure. **D. The domain organization of T1154.** The top-ranking model generated for this target is submitted by Yang-server. **E. Relationship between the baseline Neff values and the best ICS values generated for the interdomain category.** The horizontal green dotted line represents the mean best-ICS value (0.64), and the vertical red dotted line represents the mean Neff value (2117). The green zone represents the area where lower Neff values are associated with higher ICS values, while the red zone represents the area where higher Neff values are associated with lower ICS values. Neff values are depicted on the x-axis using a logarithmic scale to emphasize distinctions among low Neff values. The interdomain targets from the same structure are colored uniformly. Data point markers are scaled to their assembly size with larger complexes shown as larger circles. The depicted Neff data are tabulated in Table 6.

**Table 6.**
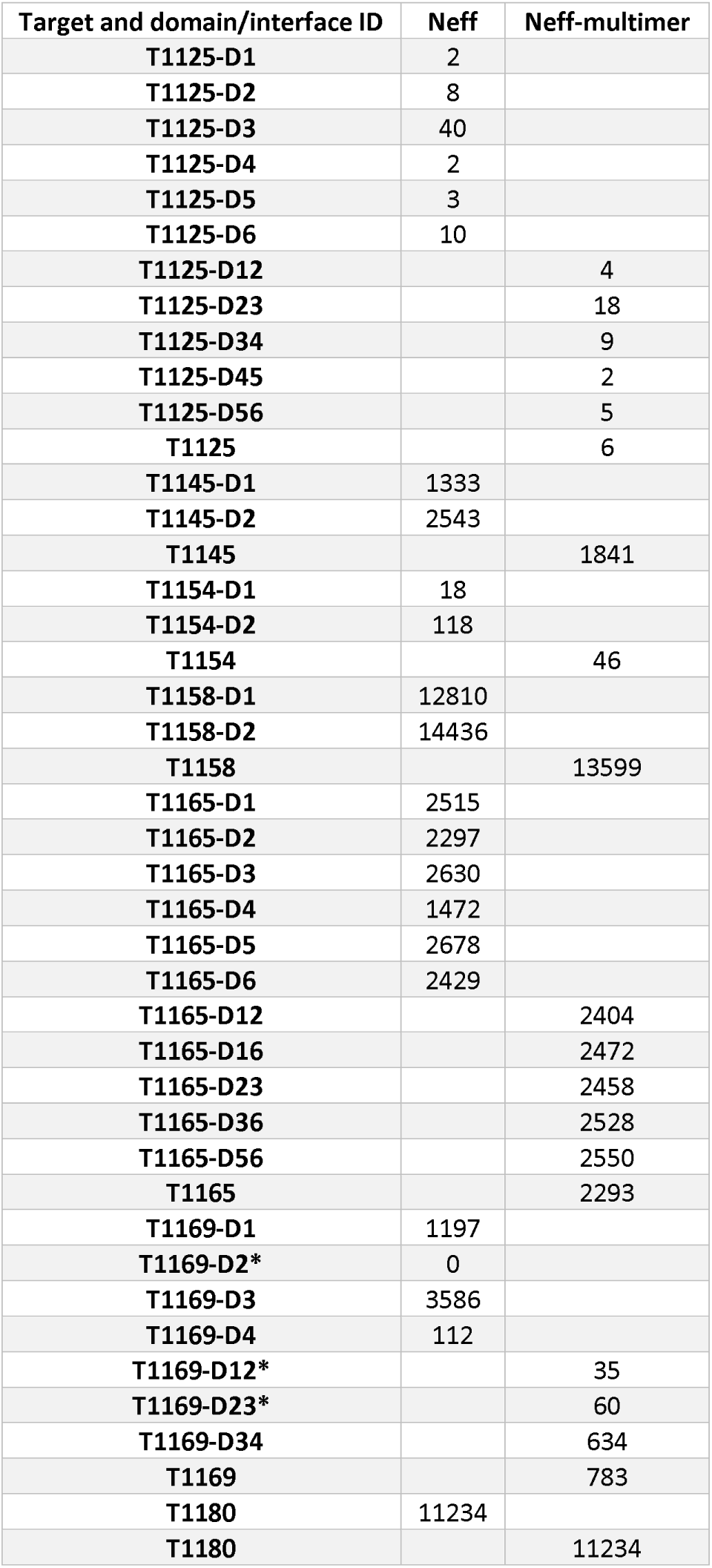
Interdomain Neff-multimer and domain Neff values calculated for the MSA obtained by standard AF2 sampling. D denotes domains. Neff-multimer calculation is performed as described in Methods.

To investigate the impact of MSAs on prediction success, we plotted the best ICS for each domain combination versus the depth of their combined MSAs (Neff, see Methods) (Figure 7E). Similar to the assembly category, we divided the plot into four zones based on the mean ICS and Neff values and focused on the analysis of targets in green and red zones. The red zone includes T1165 targets: T1165, T1165-D16, T1165-D56. The green zone includes seven targets: T1125-D23, T1125-D45, T1145, T1154, T1169, T1169-D23, T1169-D34.

Among the green zone targets, T1125-D23, T1125-D45, T1154, T1169, and T1169-D23 have extremely low Neff values. As an outcome, in the case of T1125 and in T1169, the lack of MSA information hindered achieving the required domain compaction. T1125 is a gp96RNAP enzyme found in Thermus phage. Among the six domains of T1125, only D3 has a moderate Neff value, while the remaining domains have extremely low Neff values. Even so, three interfaces of this target, namely T1125-D23, T1125-D45, and T1125-D56 were successfully predicted. However, having these three interfaces modeled properly was not enough to model the whole system, since the wrongly predicted D12 and D34 interfaces hindered the necessary closing of the N-terminal region of this protein upon its C-terminal region (Figure 7B). T1169 is a mosquito salivary gland surface protein 1 with more than 3000 residues. It consists of four domains and three qualified interfaces (Table 5). Despite the very low Neff values for its D2 and D4 two domains, D23 and D34 could be predicted successfully. As in T1125, in this case too, accurate prediction of D12 was needed for the proper folding of the protein, which could not be achieved by anyone (Figure 7C). The final extremely low Neff case, T1154, is an archaeal protein with two domains and one interface for evaluation. This Y-shaped target is mainly made of beta sheets. Despite having an extremely low Neff value, it was surprisingly well predicted by many predictors (Figure 7D). This tells us that this target somehow falls into the “learned structural space” of the participating AI models.

The final interdomain target worth mentioning is T1165, which is a modified construct of the native ubiquitin-protein ligase TOM1 from Saccharomyces cerevisiae. This protein is composed of more than 3000 amino acids, made of six domains and five qualified interfaces. Among these five interfaces, four could be successfully predicted, however the models for three of them, i.e., T1165, T1165-D16, T1165-D56, fell into the red zone. The interfaces falling into the red zone despite a high Neff suggests that the large size of the protein may be a limiting factor in these cases. Additionally, the only mispredicted interface was D16 (Figure S9). The D1 and D6 domains are made of stacking of several alpha helices as in TPR repeat proteins. Predicting the proper stacking of these alpha helices posed a challenge, which resulted in the misprediction of the D1-D6 interface. As an outcome, the 894 Å² D1-D6 interface was predicted to be as small as 25.5 Å² by the top predictor of this target (Figure S9). This highlights the challenge in predicting the overall fold of the alpha helical repeat proteins.

## CONCLUSIONS AND PERSPECTIVES

The CASP15 assembly category witnessed unprecedented advancements in assembly modeling, achieving an impressive success rate of 90% compared to previous rounds. This remarkable progress can be attributed to the improved performance of AF2M, achieved through the utilization of custom-built MSAs and the optimization of sampling parameters. In order to explore the potential of combining MSA-and sampling-focused approaches, we conducted a detailed analysis on a per-case basis, comparing the models generated by Zheng and Wallner based on their best ICS scores (Figure S10). The analysis revealed that Wallner demonstrated excellent performance in cases involving mutation-induced, host-pathogen, and nanobody-antigen complexes, while Zheng excelled in modeling large assemblies (Figure S10). Such a combination, though, would still be limited in modeling antibody-antigen complexes or constructing very large assemblies. To this end, two recently proposed methodologies can come to rescue. The first one, AlphaLink, incorporates experimental distances obtained from cross-linking mass spectrometry to guide AF2M-based complex formation^80,81^. The second one, RoseTTAFold2, leverages the strengths of both RoseTTAFold and AF2 and has shown scalability in modeling large complexes^82^. Additionally, the success of CASP15 has motivated DeepMind to release an updated version of AF2, known as AF2-v2.3.0^83^. This update has successfully addressed sampling challenges encountered in standard AF2M for modeling host-pathogen interactions (H1129), nanobody-antigen complexes (H1144), and large assemblies (H1111 and H1114) (Figure S11).

### Notes to the CASP16 Assembly Assessor

In the CASP15 assembly category, we followed the conventional CASP difficulty classification. However, based on our assessment, we recommend updating the difficulty classification in the next round to consider both MSA depth and available structural models. Additionally, we found that ICS was an effective metric for discriminating between the top-ranking groups. However, upon further analysis, we identified a limitation in ICS scores for very large complexes with multiple interfaces, where the scores were inflated due to the dominance of larger interfaces (Figure S12). To address this issue, we propose either upweighting the smaller interfaces, or evaluating each interface separately. Furthermore, in future rounds, it would be beneficial to incorporate target resolution information into the accuracy calculation. In the current round, we observed that targets with >3Å resolution consistently exhibited ICS values ranging from 0.6 to 0.8 (Figure S13). A systematic analysis is necessary to determine whether this lower ICS range corresponds to experimental error or other factors. Lastly, as proposed in the CASP13 Assembly Assessment paper, we suggest introducing an initial phase in the next competition round where assembly targets are provided without revealing the stoichiometry information^19^. This would present an additional challenge and encourage novel approaches in assembling protein complexes.

## CONFLICT OF INTEREST

The authors declare no conflicts of interest.

## DATA AVAILABILITY STATEMENT

The raw scoring data can be accessed at: https://predictioncenter.org/casp15/multimer_results.cgi

The processed scoring data with the accompanying scripts to generate our analysis plots can be found at: https://github.com/CSB-KaracaLab/CASP15_Assembly

## ACKNOWLEDGEMENTS

We would like to sincerely thank Claudio Mirabello from the National Bioinformatics

Infrastructure Sweden at SciLifeLab for providing us with the naïve AF2 models, along with their MSA and Neff parameters. We want to thank Gabriel Studer, the EMA assessor, for developing the approach that helped us overcome our chain mapping challenges. Our appreciation extends to all the experimental groups and predictors who actively participated in CASP15. Special acknowledgements go to the CASP organizers for their constant support throughout the competition. We also would like to wholeheartedly thank Shoshana Wodak and Marck F. Lensink for their close collaboration and invaluable assistance during the assessment.

## SUPPORTING INFORMATION

Additional supporting information is submitted with the manuscript.

## FUNDING INFORMATION

Ezgi Karaca and Burcu Ozden were supported by the EMBO Installation Grant (grant number 4421). Andriy Kryshtafovych was supported by the US National Institute of General Medical Sciences (NIGMS/NIH) grant R01GM100482.

## Notes

https://predictioncenter.org/casp15/multimer_results.cgi

### Competing Interest Statement

The authors have declared no competing interest.

### Summary of Updates

Figure 2 updated and some text clarifications are added.

## REFERENCES

1. Burley SK, Berman HM, Kleywegt GJ, Markley JL, Nakamura H, Velankar S. Protein Data Bank (PDB): The Single Global Macromolecular Structure Archive. In: ; 2017:627–641. doi:10.1007/978-1-4939-7000-1_26

2. Rout MP, Sali A. Principles for Integrative Structural Biology Studies. Cell. 2019;177(6):1384–1403. doi:10.1016/j.cell.2019.05.016

3. Akdel M, Pires DE V., Pardo EP, et al. A structural biology community assessment of AlphaFold2 applications. Nat Struct Mol Biol. 2022;29(11):1056–1067. doi:10.1038/s41594-022-00849-w

4. Humphreys IR, Pei J, Baek M, et al. Computed structures of core eukaryotic protein complexes. Science (1979). 2021;374(6573). doi:10.1126/science.abm4805

5. Varadi M, Bordin N, Orengo C, Velankar S. The opportunities and challenges posed by the new generation of deep learning-based protein structure predictors. Curr Opin Struct Biol. 2023;79:102543. doi:10.1016/j.sbi.2023.102543

6. Graziadei A, Rappsilber J. Leveraging crosslinking mass spectrometry in structural and cell biology. Structure. 2022;30(1):37–54. doi:10.1016/j.str.2021.11.007

7. Beton JG, Cragnolini T, Kaleel M, Mulvaney T, Sweeney A, Topf M. Integrating model simulation tools and <scp>cryo-electron</scp> microscopy. WIREs Computational Molecular Science. 2023;13(3). doi:10.1002/wcms.1642

8. Hopf TA, Schärfe CPI, Rodrigues JPGLM, et al. Sequence co-evolution gives 3D contacts and structures of protein complexes. Elife. 2014;3. doi:10.7554/eLife.03430

9. Andreani J, Quignot C, Guerois R. Structural prediction of protein interactions and docking using conservation and coevolution. WIREs Computational Molecular Science. 2020;10(6). doi:10.1002/wcms.1470

10. Koehler Leman J, Künze G. Recent Advances in NMR Protein Structure Prediction with ROSETTA. Int J Mol Sci. 2023;24(9):7835. doi:10.3390/ijms24097835

11. Karaca E, Bonvin AMJJ. Advances in integrative modeling of biomolecular complexes. Methods. 2013;59(3):372–381. doi:10.1016/j.ymeth.2012.12.004

12. Saltzberg D, Greenberg CH, Viswanath S, et al. Modeling Biological Complexes Using Integrative Modeling Platform. In: ; 2019:353–377. doi:10.1007/978-1-4939-9608-7_15

13. Pozzati G, Zhu W, Bassot C, Lamb J, Kundrotas P, Elofsson A. Limits and potential of combined folding and docking. Bioinformatics. 2022;38(4):954–961. doi:10.1093/bioinformatics/btab760

14. Baek M, DiMaio F, Anishchenko I, et al. Accurate prediction of protein structures and interactions using a three-track neural network. Science (2021). 2021;373(6557):871-876. doi:10.1126/science.abj8754

15. Wodak SJ, Vajda S, Lensink MF, Kozakov D, Bates PA. Critical Assessment of Methods for Predicting the 3D Structure of Proteins and Protein Complexes. Annu Rev Biophys. 2023;52(1):183–206. doi:10.1146/annurev-biophys-102622-084607

16. Janin J, Henrick K, Moult J, et al. CAPRI: A Critical Assessment of PRedicted Interactions. Proteins: Structure, Function, and Genetics. 2003;52(1):2–9. doi:10.1002/prot.10381

17. Lensink MF, Velankar S, Kryshtafovych A, et al. Prediction of homoprotein and heteroprotein complexes by protein docking and template-based modeling: A CASP-CAPRI experiment. Proteins: Structure, Function, and Bioinformatics. 2016;84(S1):323–348. doi:10.1002/prot.25007

18. Lafita A, Bliven S, Kryshtafovych A, et al. Assessment of protein assembly prediction in CASP12. Proteins: Structure, Function and Bioinformatics. 2018;86:247–256. doi:10.1002/prot.25408

19. Guzenko D, Lafita A, Monastyrskyy B, Kryshtafovych A, Duarte JM. Assessment of protein assembly prediction in CASP13. Proteins: Structure, Function and Bioinformatics. 2019;87(12):1190–1199. doi:10.1002/prot.25795

20. Ozden B, Kryshtafovych A, Karaca E. Assessment of the CASP14 assembly predictions. Proteins: Structure, Function and Bioinformatics. 2021;89(12):1787–1799. doi:10.1002/prot.26199

21. Lensink MF, Velankar S, Kryshtafovych A, et al. Prediction of homoprotein and heteroprotein complexes by protein docking and template-based modeling: A CASP-CAPRI experiment. Proteins: Structure, Function, and Bioinformatics. 2016;84(S1):323–348. doi:10.1002/prot.25007

22. Lensink MF, Brysbaert G, Mauri T, et al. Prediction of protein assemblies, the next frontier: The <scp>CASP14-CAPRI</scp> experiment. Proteins: Structure, Function, and Bioinformatics. 2021;89(12):1800-1823. doi:10.1002/prot.26222

23. Lensink MF, Brysbaert G, Nadzirin N, et al. Blind prediction of homo-and hetero-protein complexes: The CASP13-CAPRI experiment. Proteins: Structure, Function, and Bioinformatics. 2019;87(12):1200–1221. doi:10.1002/prot.25838

24. Lensink MF, Velankar S, Baek M, Heo L, Seok C, Wodak SJ. The challenge of modeling protein assemblies: the CASP12-CAPRI experiment. Proteins: Structure, Function, and Bioinformatics. 2018;86:257–273. doi:10.1002/prot.25419

25. Soni N, Madhusudhan MS. Computational modeling of protein assemblies. Curr Opin Struct Biol. 2017;44:179–189. doi:10.1016/j.sbi.2017.04.006

26. Jumper J, Evans R, Pritzel A, et al. Highly accurate protein structure prediction with AlphaFold. Nature. 2021;596(7873):583-589. doi:10.1038/s41586-021-03819-2

27. Kryshtafovych A, Schwede T, Topf M, Fidelis K, Moult J. Critical assessment of methods of protein structure prediction (CASP)—Round <scp>XIV</scp>. Proteins: Structure, Function, and Bioinformatics. 2021;89(12):1607-1617. doi:10.1002/prot.26237

28. Varadi M, Anyango S, Deshpande M, et al. AlphaFold Protein Structure Database: massively expanding the structural coverage of protein-sequence space with high-accuracy models. Nucleic Acids Res. 2022;50(D1):D439–D444. doi:10.1093/nar/gkab1061

29. Callaway E. ‘The entire protein universe’: AI predicts shape of nearly every known protein. Nature. 2022;608(7921):15-16. doi:10.1038/d41586-022-02083-2

30. Tsaban T, Varga JK, Avraham O, Ben-Aharon Z, Khramushin A, Schueler-Furman O. Harnessing protein folding neural networks for peptide–protein docking. Nat Commun. 2022;13(1):176. doi:10.1038/s41467-021-27838-9

31. Ko J, Lee J. Can AlphaFold2 predict protein-peptide complex structures accurately? biorxiv. Published online July 28, 2021.

32. Mirdita M, Schütze K, Moriwaki Y, Heo L, Ovchinnikov S, Steinegger M. ColabFold: making protein folding accessible to all. Nat Methods. 2022;19(6):679–682. doi:10.1038/s41592-022-01488-1

33. Evans R, O’neill M, Pritzel A, et al. Protein complex prediction with AlphaFold-Multimer. Published online 2022. doi:10.1101/2021.10.04.463034

34. Bryant P, Pozzati G, Elofsson A. Improved prediction of protein-protein interactions using AlphaFold2. Nat Commun. 2022;13(1). doi:10.1038/s41467-022-28865-w

35. Yin R, Feng BY, Varshney A, Pierce BG. Benchmarking <scp>AlphaFold</scp> for protein complex modeling reveals accuracy determinants. Protein Science. 2022;31(8). doi:10.1002/pro.4379

36. Gao M, Nakajima An D, Parks JM, Skolnick J. AF2Complex predicts direct physical interactions in multimeric proteins with deep learning. Nat Commun. 2022;13(1):1744. doi:10.1038/s41467-022-29394-2

37. Elofsson A. Progress at protein structure prediction, as seen in CASP15. Curr Opin Struct Biol. 2023;80:102594. doi:10.1016/j.sbi.2023.102594

38. Pagès G, Grudinin S. AnAnaS: Software for Analytical Analysis of Symmetries in Protein Structures. In: ; 2020:245–257. doi:10.1007/978-1-0716-0708-4_14

39. Duarte JM, Srebniak A, Schärer MA, Capitani G. Protein Interface Classification by Evolutionary Analysis.; 2012. http://www.biomedcentral.com/1471-2105/13/334

40. Krissinel E, Henrick K. Inference of Macromolecular Assemblies from Crystalline State. J Mol Biol. 2007;372(3):774–797. doi:10.1016/j.jmb.2007.05.022

41. Protein interfaces, surfaces and assemblies. service PISA at the European Bioinformatics Institute. Accessed August 31, 2023. http://www.ebi.ac.uk/pdbe/prot_int/pistart.html

42. Zimmermann L, Stephens A, Nam SZ, et al. A Completely Reimplemented MPI Bioinformatics Toolkit with a New HHpred Server at its Core. J Mol Biol. 2018;430(15):2237–2243. doi:10.1016/j.jmb.2017.12.007

43. Gabler F, Nam S, Till S, et al. Protein Sequence Analysis Using the MPI Bioinformatics Toolkit. Curr Protoc Bioinformatics. 2020;72(1). doi:10.1002/cpbi.108

44. Studer G, Tauriello G, Schwede T. All about complexes. Proteins. 2023; Prot-00157-2023, final review, this issue

45. Xu J, Zhang Y. How significant is a protein structure similarity with TM-score = 0.5? Bioinformatics. 2010;26(7):889–895. doi:10.1093/bioinformatics/btq066

46. Mariani V, Biasini M, Barbato A, Schwede T. lDDT: a local superposition-free score for comparing protein structures and models using distance difference tests. Bioinformatics. 2013;29(21):2722–2728. doi:10.1093/bioinformatics/btt473

47. Critical Assessment Of Techniques For Protein Structure Prediction Abstract Book: Fifteenth Round.; 2022. Accessed June 18, 2023. https://predictioncenter.org/casp15/doc/CASP15_Abstracts.pdf

48. Mirabello C. NBIS-AF2-M Results. Accessed June 18, 2023. http://duffman.it.liu.se/casp15/

49. Johnson LS, Eddy SR, Portugaly E. Hidden Markov Model Speed Heuristic and Iterative HMM Search Procedure.; 2010. http://ekhidna.biocenter.

50. Wu T, Hou J, Adhikari B, Cheng J. Analysis of several key factors influencing deep learning-based inter-residue contact prediction. Bioinformatics. 2020;36(4):1091–1098. doi:10.1093/bioinformatics/btz679

51. Bryant P, Pozzati G, Zhu W, Shenoy A, Kundrotas P, Elofsson A. Predicting the structure of large protein complexes using AlphaFold and Monte Carlo tree search. doi:10.1101/2022.03.12.484089

52. Vangone A, Spinelli R, Scarano V, Cavallo L, Oliva R. COCOMAPS: a web application to analyze and visualize contacts at the interface of biomolecular complexes. Bioinformatics. 2011;27(20):2915–2916. doi:10.1093/bioinformatics/btr484

53. Bertoni M, Kiefer F, Biasini M, Bordoli L, Schwede T. Modeling protein quaternary structure of homo-and hetero-oligomers beyond binary interactions by homology. Sci Rep. 2017;7(1):10480. doi:10.1038/s41598-017-09654-8

54. Xu Q, Canutescu AA, Wang G, Shapovalov M, Obradovic Z, Dunbrack RL. Statistical Analysis of Interface Similarity in Crystals of Homologous Proteins. J Mol Biol. 2008;381(2):487–507. doi:10.1016/j.jmb.2008.06.002

55. Hunter JD. Matplotlib: A 2D graphics environment. Comput Sci Eng. 2007;9(3):90–95.

56. Waskom ML. seaborn: statistical data visualization. J Open Source Softw. 2021;6(60):3021. doi:10.21105/joss.03021

57. Harris CR, Millman KJ, der Walt SJ van, et al. Array programming with NumPy. Nature. 2020;585(7825):357-362. doi:10.1038/s41586-020-2649-2

58. Van Rossum G, Drake FL. Python 3 Reference Manual. CreateSpace; 2009.

59. McKinney W, others. Data structures for statistical computing in python. In: Proceedings of the 9th Python in Science Conference. Vol 445. ; 2010:51-56.

60. Schrödinger LLC, DeLano W. PyMOL 2.5.2. http://www.pymol.org/pymol

61. Raasakka A, Myllykoski M, Laulumaa S, et al. Determinants of ligand binding and catalytic activity in the myelin enzyme 21,31-cyclic nucleotide 31-phosphodiesterase. Sci Rep. 2015;5(1):16520. doi:10.1038/srep16520

62. Yagi S, Padhi AK, Vucinic J, et al. Seven Amino Acid Types Suffice to Create the Core Fold of RNA Polymerase. J Am Chem Soc. 2021;143(39):15998–16006. doi:10.1021/jacs.1c05367

63. Zheng W, Wuyun Q, Freddolino PL, Zhang Y. Integrating deep learning, threading alignments, and a <scp>multi-MSA</scp> strategy for high-quality protein monomer and complex structure prediction in <scp>CASP15</scp>. Proteins: Structure, Function, and Bioinformatics. Published online August 31, 2023. doi:10.1002/prot.26585

64. Remmert M, Biegert A, Hauser A, Söding J. HHblits: lightning-fast iterative protein sequence searching by HMM-HMM alignment. Nat Methods. 2012;9(2):173–175. doi:10.1038/nmeth.1818

65. Potter SC, Luciani A, Eddy SR, Park Y, Lopez R, Finn RD. HMMER web server: 2018 update. Nucleic Acids Res. 2018;46(W1):W200-W204. doi:10.1093/nar/gky448

66. Mirdita M, Steinegger M, Breitwieser F, Söding J, Levy Karin E. Fast and sensitive taxonomic assignment to metagenomic contigs. Bioinformatics. 2021;37(18):3029–3031. doi:10.1093/bioinformatics/btab184

67. Ritchie DW, Kemp GJL. Protein docking using spherical polar Fourier correlations. Proteins: Structure, Function, and Bioinformatics. 2000;39(2):178–194.

68. Gabb HA, Jackson RM, Sternberg MJE. Modelling protein docking using shape complementarity, electrostatics and biochemical information 1 1Edited by J. Thornton. J Mol Biol. 1997;272(1):106–120. doi:10.1006/jmbi.1997.1203

69. Ritchie DW, Grudinin S. Spherical polar Fourier assembly of protein complexes with arbitrary point group symmetry. J Appl Crystallogr. 2016;49(1):158–167. doi:10.1107/S1600576715022931

70. Kliment O, Valančauskas L, Dapkūnas J, Venclovas Č. Prediction of protein assemblies by structure sampling followed by interface-focused scoring. biorxiv. Published online March 8, 2023.

71. Olechnovič K, Valančauskas L, Dapkūnas J, Venclovas Č. Prediction of protein assemblies by structure sampling followed by interface-focused scoring. Proteins: Structure, Function, and Bioinformatics. Published online August 14, 2023. doi:10.1002/prot.26569

72. Wallner B. AFsample: Improving Multimer Prediction with AlphaFold using Aggressive Sampling. Published online 2023. doi:10.1101/2022.12.20.521205

73. Söding J. Protein homology detection by HMM–HMM comparison. Bioinformatics. 2005;21(7):951–960. doi:10.1093/bioinformatics/bti125

74. Altschul S. Gapped BLAST and PSI-BLAST: a new generation of protein database search programs. Nucleic Acids Res. 1997;25(17):3389–3402. doi:10.1093/nar/25.17.3389

75. Oda T, Lim K, Tomii K. Simple adjustment of the sequence weight algorithm remarkably enhances PSI-BLAST performance. BMC Bioinformatics. 2017;18(1):288. doi:10.1186/s12859-017-1686-9

76. Mori H, Ishikawa H, Higashi K, Kato Y, Ebisuzaki T, Kurokawa K. PZLAST: an ultra-fast amino acid sequence similarity search server against public metagenomes. Bioinformatics. 2021;37(21):3944–3946. doi:10.1093/bioinformatics/btab492

77. Kryshtafovych A, Antczak M, Szachniuk M, et al. New prediction categories in CASP15. Published online 2023. doi:10.22541/au.168371643.39402261/v1

78. Rigden D, Simpkin A, Mesdaghi S, et al. Tertiary structure assessment at CASP15. Published online 2023. doi:10.22541/au.168495671.14377331/v1

79. Kryshtafovych A, Rigden D, Rigden DJ. To split or not to split: CASP15 targets and their processing into tertiary structure evaluation units. Published online March 13, 2023. doi:10.22541/au.167872023.39044035/v1

80. Stahl K, Brock O, Rappsilber J, Der SM”, Mensch S. Modelling protein complexes with crosslinking mass spectrometry and deep learning. doi:10.1101/2023.06.07.544059

81. Stahl K, Graziadei A, Dau T, Brock O, Rappsilber J. Protein structure prediction with in-cell photo-crosslinking mass spectrometry and deep learning. Nat Biotechnol. Published online 2023. doi:10.1038/s41587-023-01704-z

82. Baek M, Anishchenko I, Humphreys IR, Cong Q, Baker D, Dimaio F. Efficient and accurate prediction of protein structure using RoseTTAFold2. doi:10.1101/2023.05.24.542179

83. AlphaFold v2.3.0. Accessed June 18, 2023. https://github.com/deepmind/alphafold/blob/main/docs/technical_note_v2.3.0.md

84. Gilzer D, Schreiner M, Niemann HH. Direct interaction of a chaperone-bound type III secretion substrate with the export gate. Nat Commun. 2022;13(1):2858. doi:10.1038/s41467-022-30487-1

85. Grinter R, Kropp A, Venugopal H, et al. Structural basis for bacterial energy extraction from atmospheric hydrogen. Nature. 2023;615(7952):541-547. doi:10.1038/s41586-023-05781-7

86. van den Berg B, Silale A, Baslé A, Brandner AF, Mader SL, Khalid S. Structural basis for host recognition and superinfection exclusion by bacteriophage T5. Proceedings of the National Academy of Sciences. 2022;119(42). doi:10.1073/pnas.2211672119

87. Cuthbert BJ, Goulding CW, Hayes CS. Toxin/immunity complex for a T6SS lipase effector from E. cloacae. 10.2210/pdb7UBZ/pdb

88. Chen J, Fruhauf A, Fan C, et al. Structure of an endogenous mycobacterial MCE lipid transporter. Res Sq. Published online January 10, 2023. doi:10.21203/rs.3.rs-2412186/v1

89. Wald J, Fahrenkamp D, Goessweiner-Mohr N, et al. Mechanism of AAA+ ATPase-mediated RuvAB–Holliday junction branch migration. Nature. 2022;609(7927):630-639. doi:10.1038/s41586-022-05121-1

90. Deep A, Gu Y, Gao YQ, et al. The SMC-family Wadjet complex protects bacteria from plasmid transformation by recognition and cleavage of closed-circular DNA. Mol Cell. 2022;82(21):4145–4159.e7. doi:10.1016/j.molcel.2022.09.008

91. Delgado-Cunningham K, López T, Khatib F, Arias CF, DuBois RM. Structure of the divergent human astrovirus MLB capsid spike. Structure. 2022;30(12):1573–1581.e3. doi:10.1016/j.str.2022.10.010

92. Wu K, Moore JA, Miller MD, et al. Expanding the eukaryotic genetic code with a biosynthesized 21st amino acid. Protein Science. 2022;31(10). doi:10.1002/pro.4443

93. Bloch Y, Savvides SN. Tobacco lectin Nictaba in complex with triacetylchitotriose (CASP target). 10.2210/pdb8AD2/pdb

94. Liu S, Xia X, Calvo E, Zhou ZH. Native structure of mosquito salivary protein uncovers domains relevant to pathogen transmission. Nat Commun. 2023;14(1):899. doi:10.1038/s41467-023-36577-y

