## Supporting Information for "The Impact of AI-Based Modeling on the Accuracy of Protein Assembly Prediction: Insights from CASP15"

**Correspondence**

Ezgi Karaca

**FIGURES:**

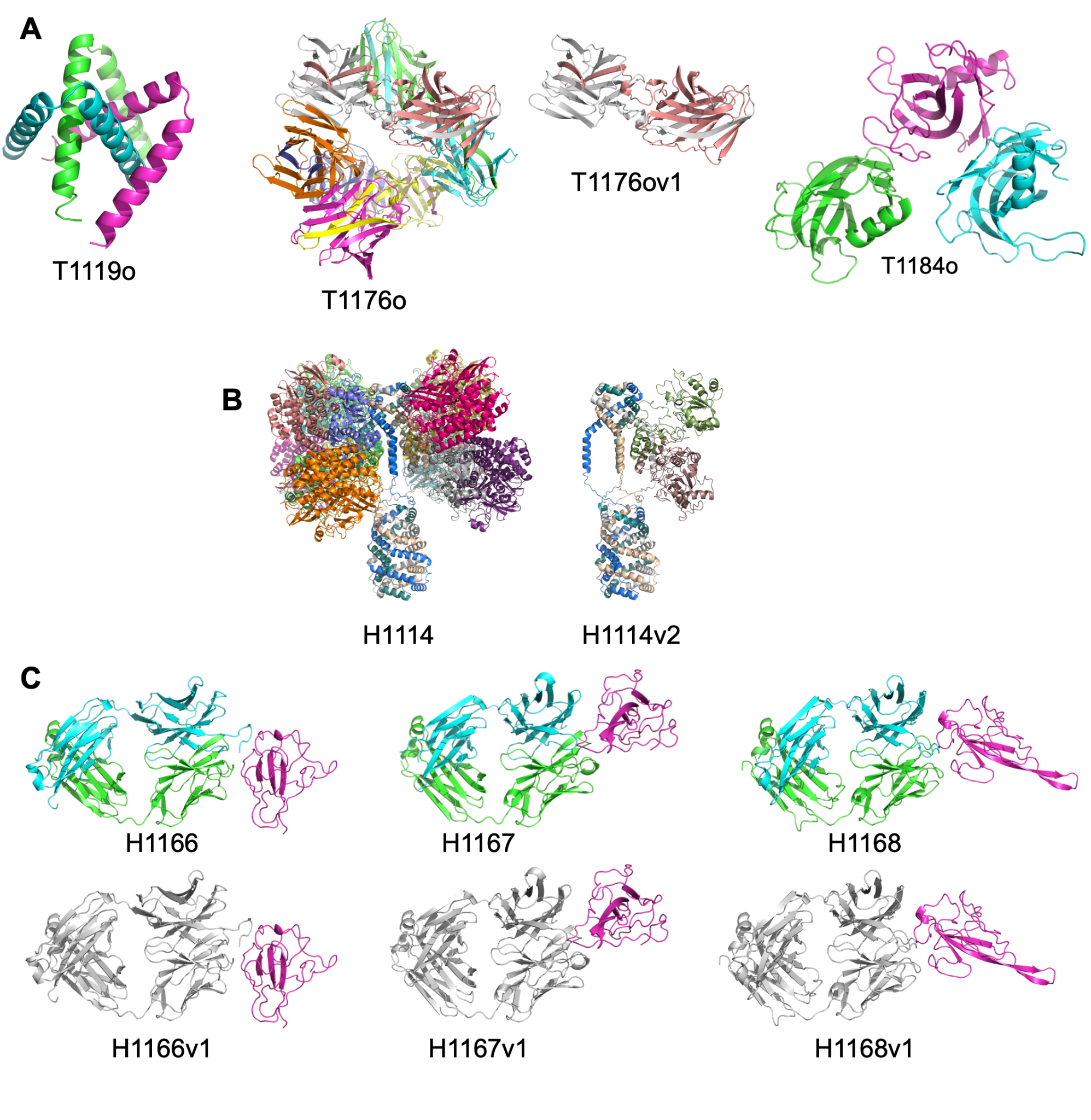

**Figure S1. Canceled and modified targets. A.** T1119o, T1176o, T1176ov1, and T1184o targets were canceled due to the inconsistencies between their assigned and predicted stoichiometries. **B.** A subcomplex of H1114 (A4B8C8) was considered as a separate target as H1114v2 (A4B2). **C.** Heavy and light chains of H1166, H1167, and H1168 were fused into one evaluation unit, leading to an updated chain organization. The updated targets were presented under target ids H1166v1, H1167v1 and H1168v1. In this figure, we colored each chain or evaluation unit differently.

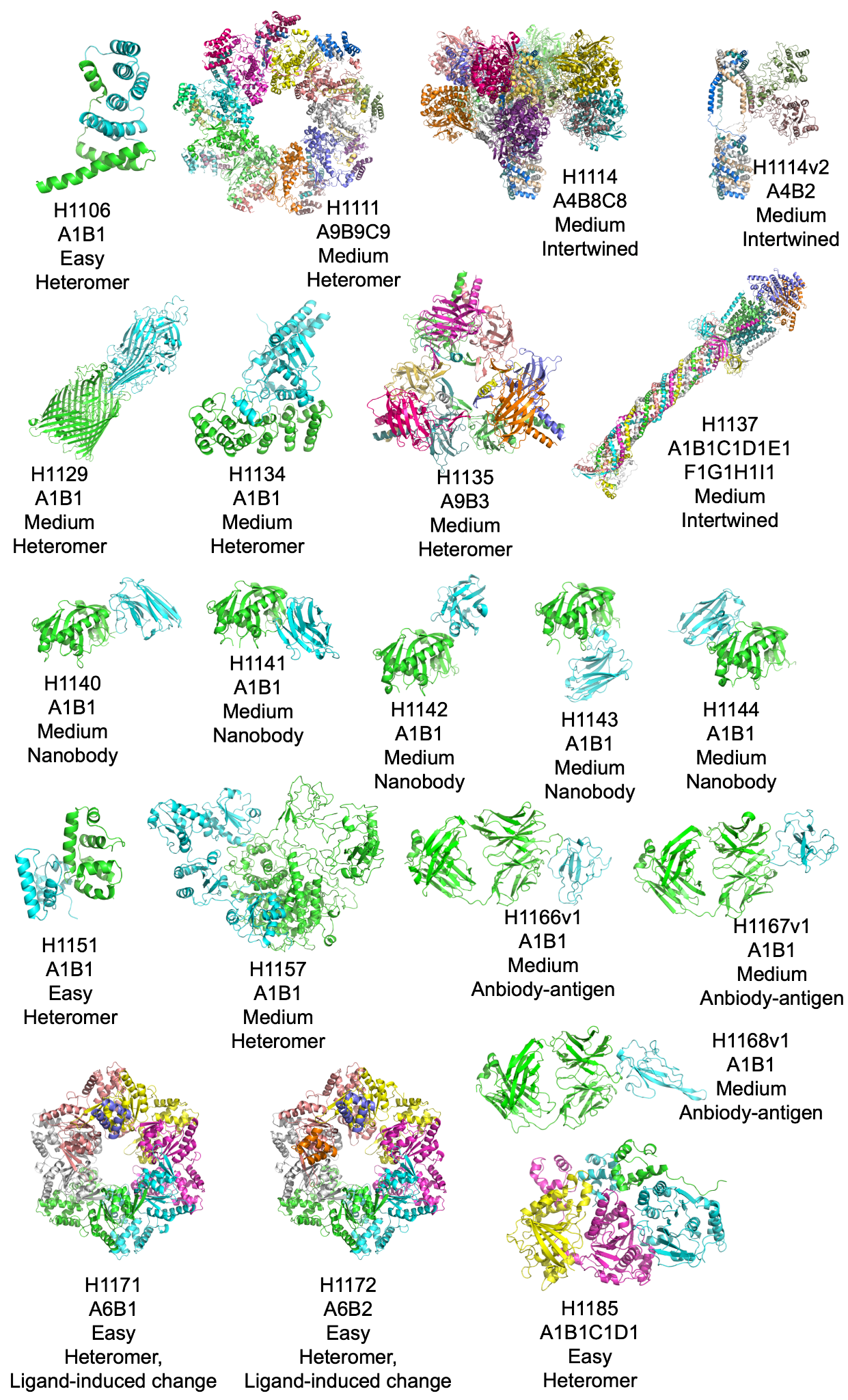

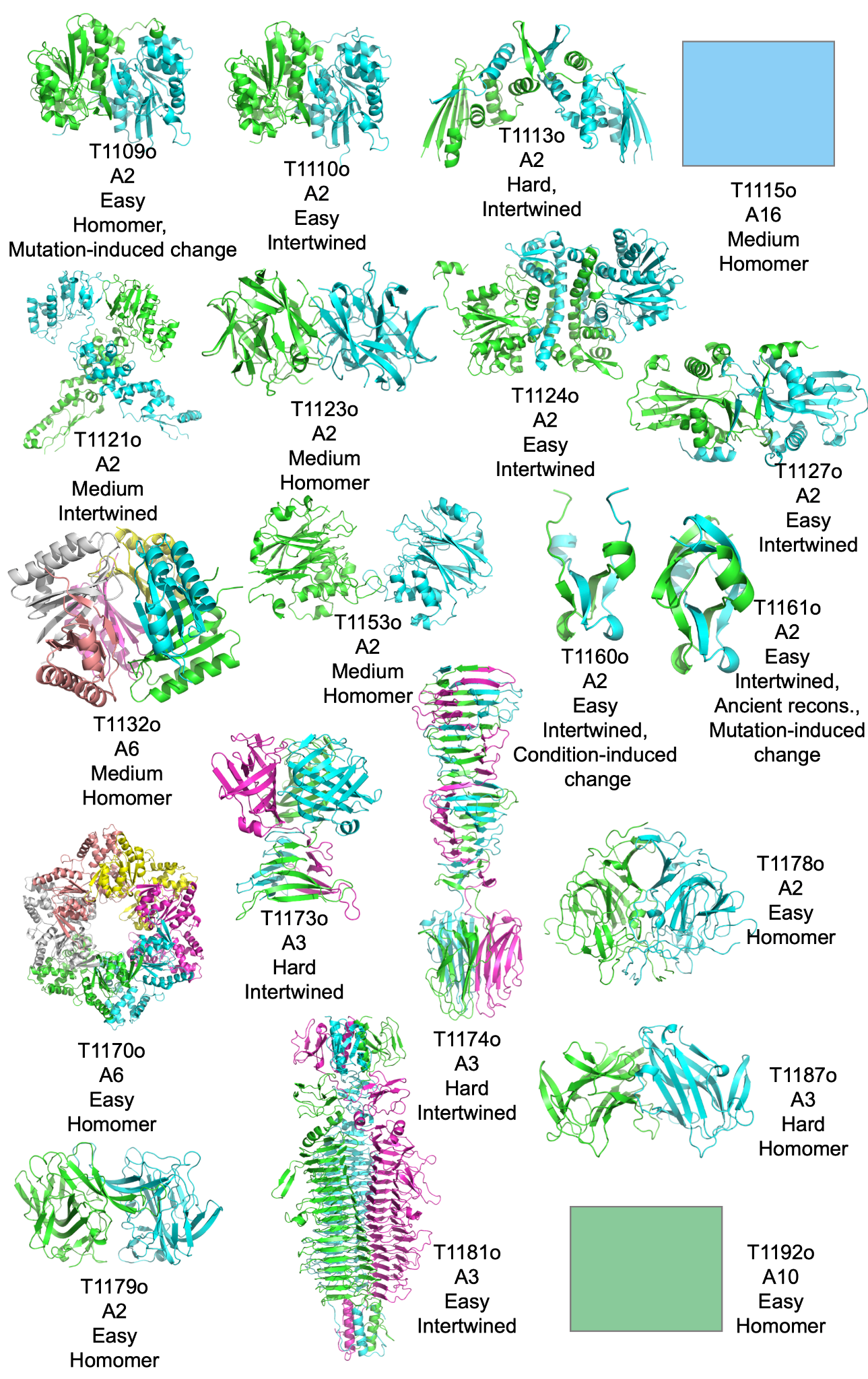

**Figure S2. CASP15 assembly targets,** together with their stoichiometries, difficulty and structural classifications. Each assembly is colored differently according to their chain ids (following the chain id coloring order of Pymol). T115o and T1192o are redacted as their structures were under embargo at the time of this publication.

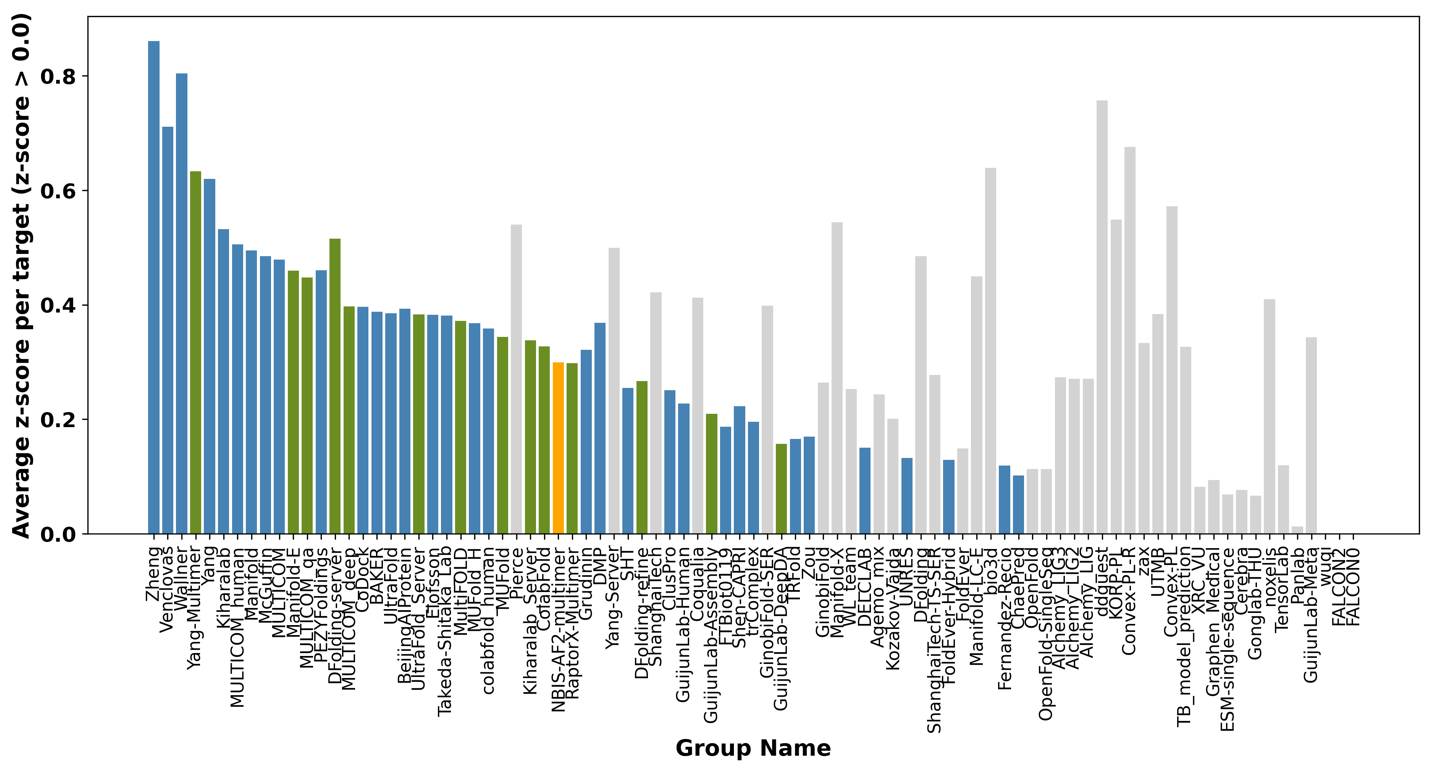

**Figure S3. The CASP15 assembly groups’ average z-scores per target**, ordered according to final ranking given in Figure 3A. The bar height represents the average z-score per target. Human groups are shown in blue, servers in green, the baseline NBIS-AF2-Multimer in orange and if any group participated less than 75% of the targets (31/41) is depicted in gray, since it may not be appropriate to make a general comment about their methodologies based on their average z-score per target.

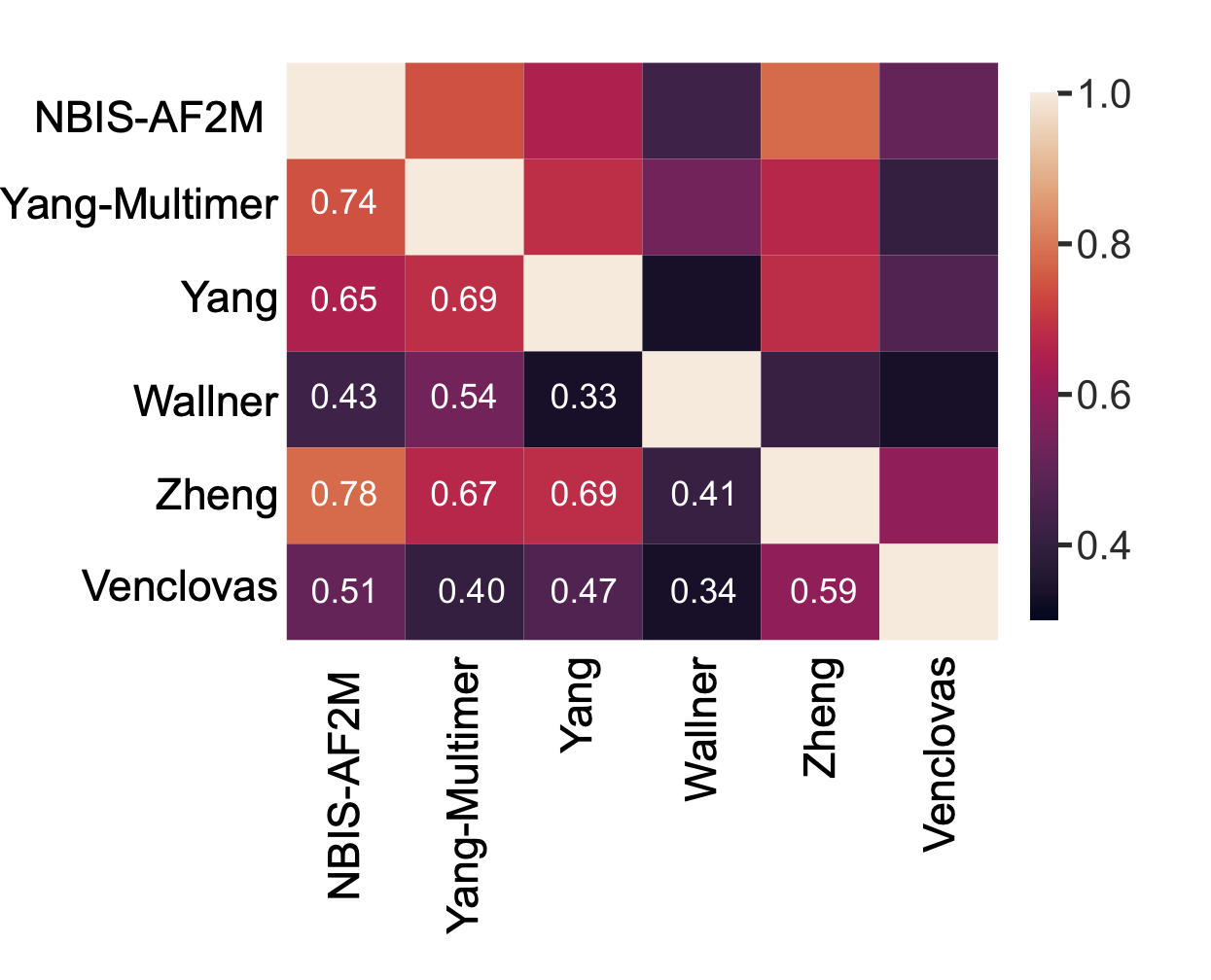

**Figure S4. Model similarities between the top-ranking groups and the baseline predictor (NBIS-AF2M).** The correlation scores were calculated over ICS values. The scores range from 0.0 to 1.0, with 1.0 indicating a perfect correlation. Light colors denote higher correlations, while dark colors denote lower ones.

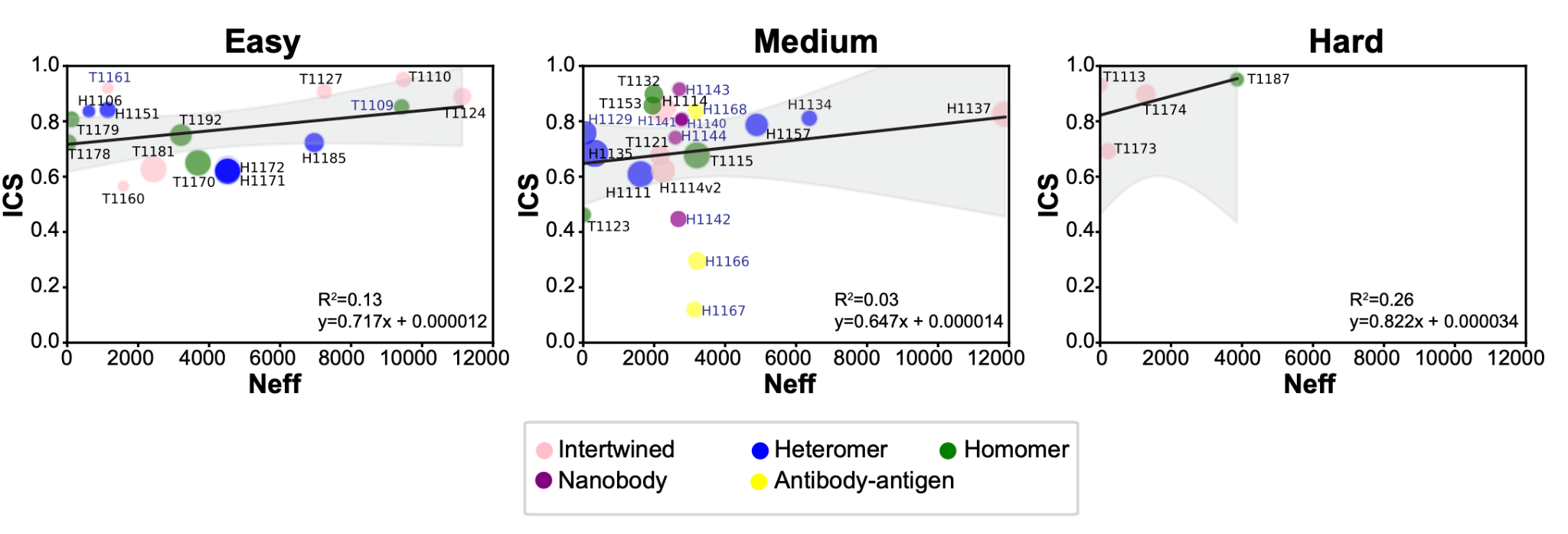

**Figure S5. Relationship between baseline multimeric Neff values and the best ICS values for each difficulty category with regression lines and 95% confidence intervals.** In all categories, the black line represents the linear regression lines and gray areas indicate the 95% confidence intervals. Markers of data points are scaled to the assembly size where larger complexes shown with larger circles. Also, markers are colored according to their structural properties (pink: intertwined, purple: nanobody, yellow: antibody-antigen, blue: heteromer, and green: homomer)

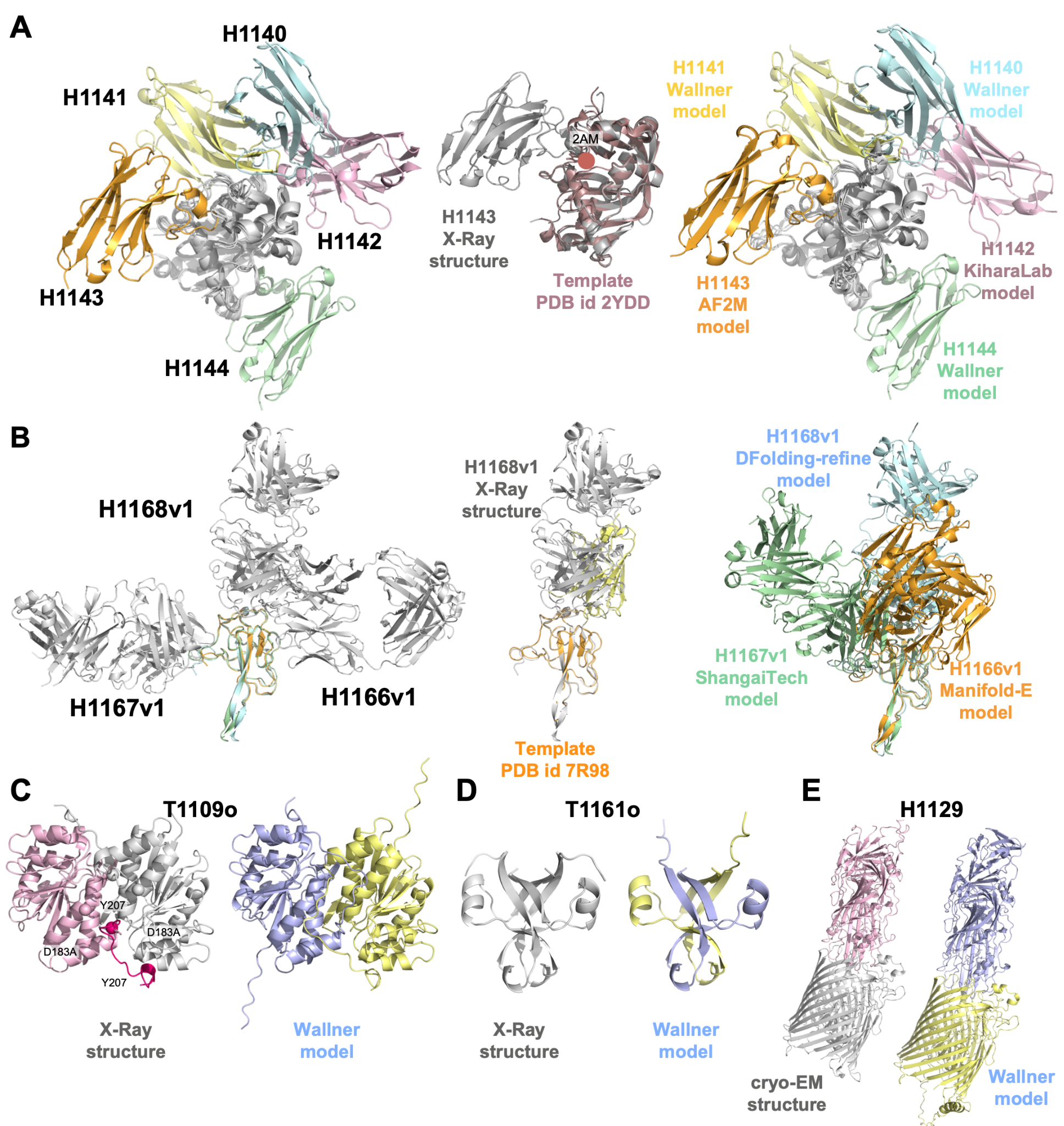

**Figure S6. Assembly target highlights where enhanced sampling makes a difference. A.** Nanobody-target complex series (H1140 - H1144), **B.** Antibody-antigen complex series (H1166v1 - H1168v1), **C.** Mutation-induced targets T1109o and **D.** T1161o, **E.** Host-pathogen target H1129. Experimental structures and their best ICS models are colored according to their subunits. For nanobody-target H1143 and antibody-antigen target H1168v1 have partial templates for their binding sites. These templates were aligned on top of the experimental structures.

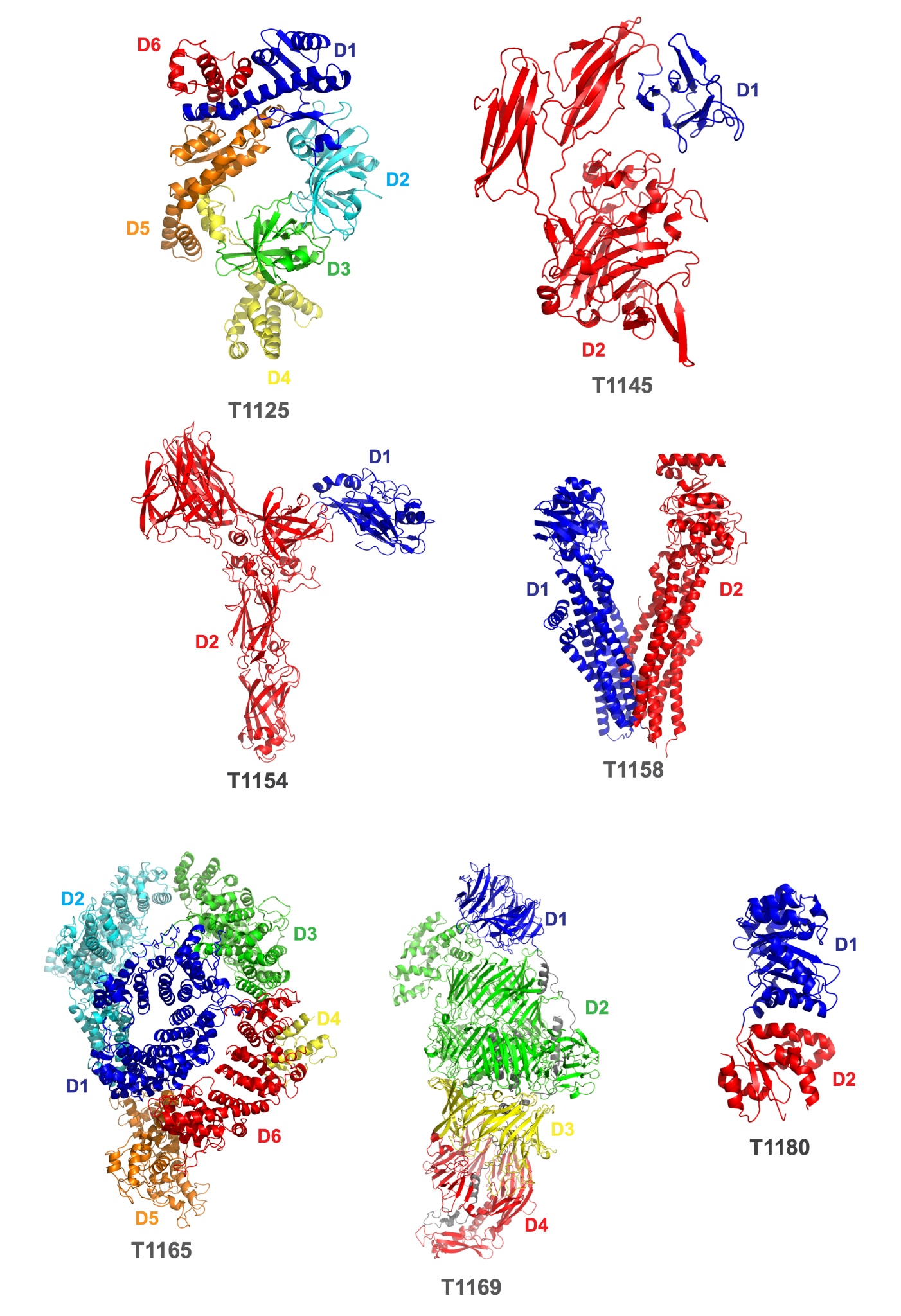

**Figure S7. Tertiary structures of interdomain targets and their evaluation units.** The structures are colored according to domain definitions.

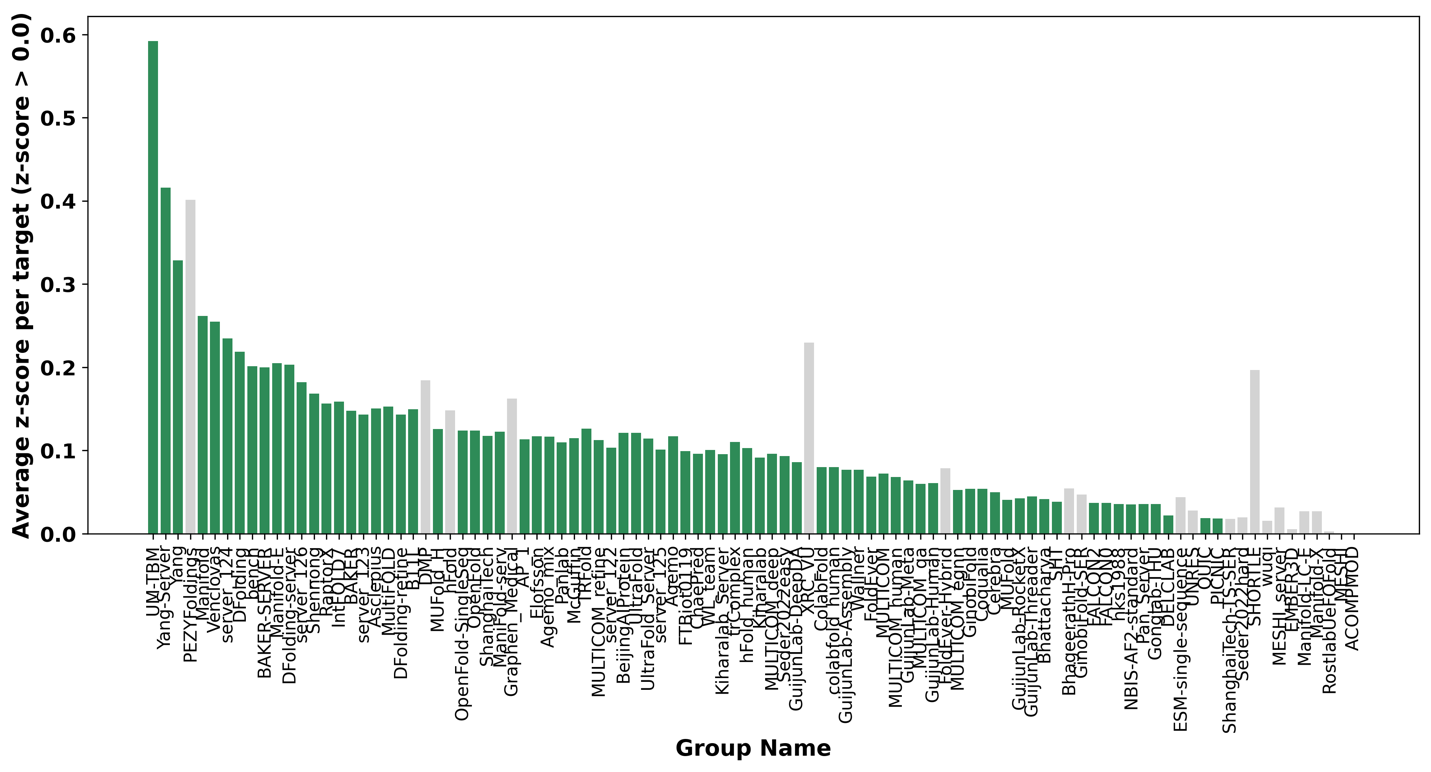

**Figure S8. Average z-scores per target of CASP15 groups in predicting interfaces of multi-domain targets.** The groups are ranked according to final ranking given in Figure 7A and any group participated less than 75% of the targets (17/22) is depicted in gray, since it may not be appropriate to make a general comment about their methodologies based on their average z-score per target.

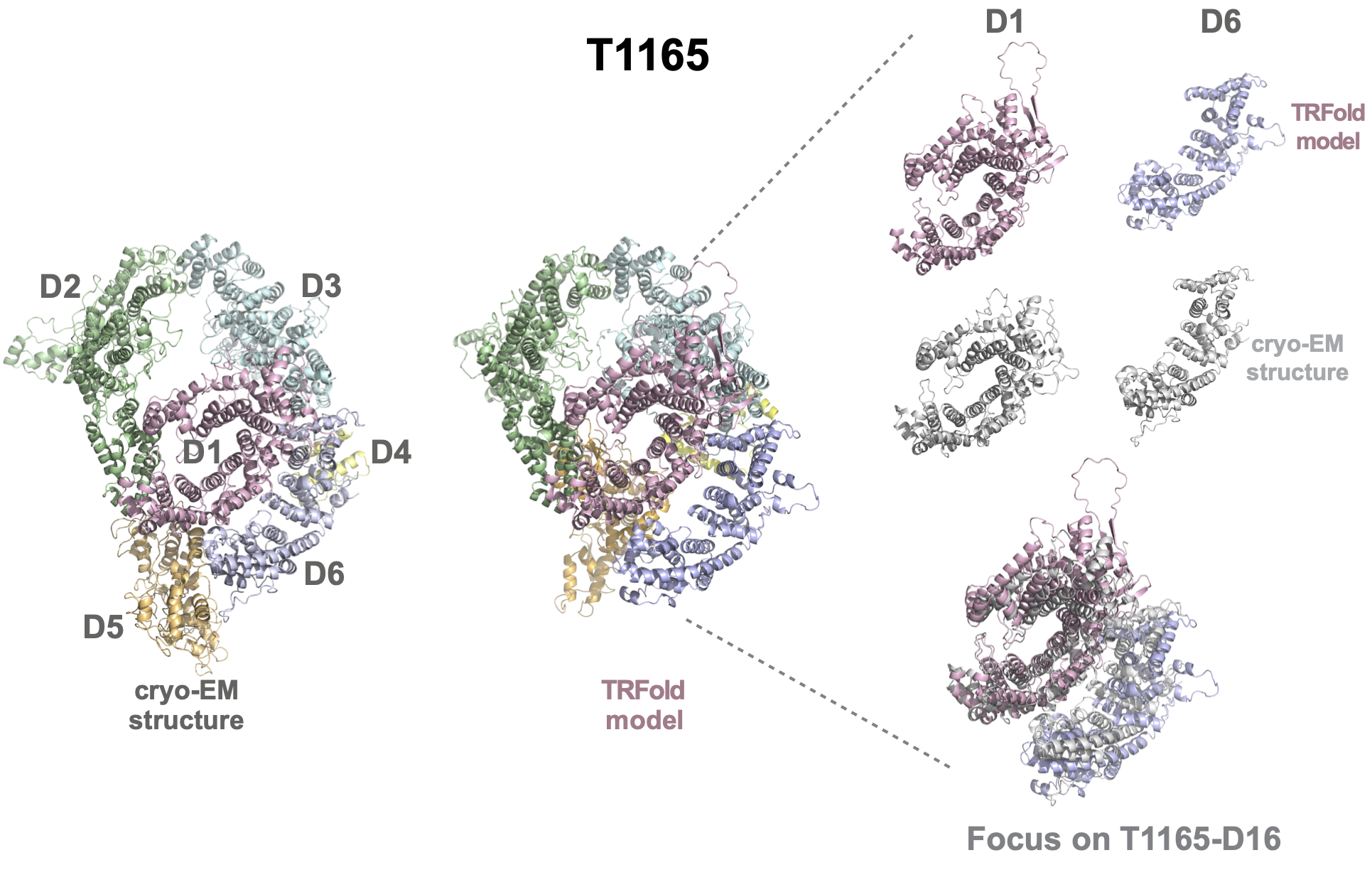
**Figure S9. The domain organization of T1165.** The top-ranking model for this target was submitted by TRFold. Among the five qualified interfaces (D12, D16, D23, D36, and D56), D16 could not be successfully modeled by any predictor. D1 and D6 reference and TRFold domain structures are compared in the inset, together with their aligned D16 interdomain complexes.

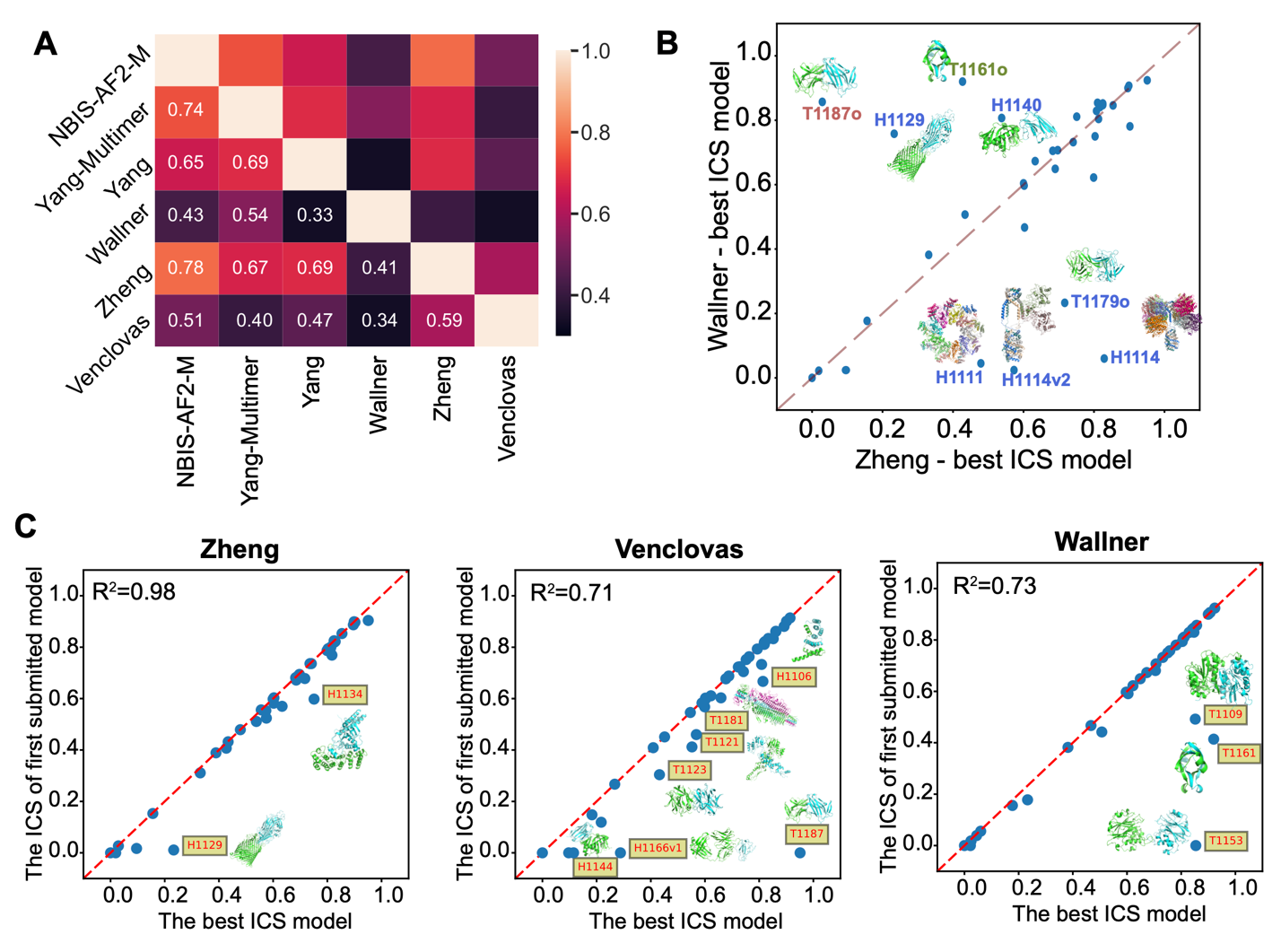

**Figure S10.** The ICS relationship between the best models submitted by Wallner and Zheng.

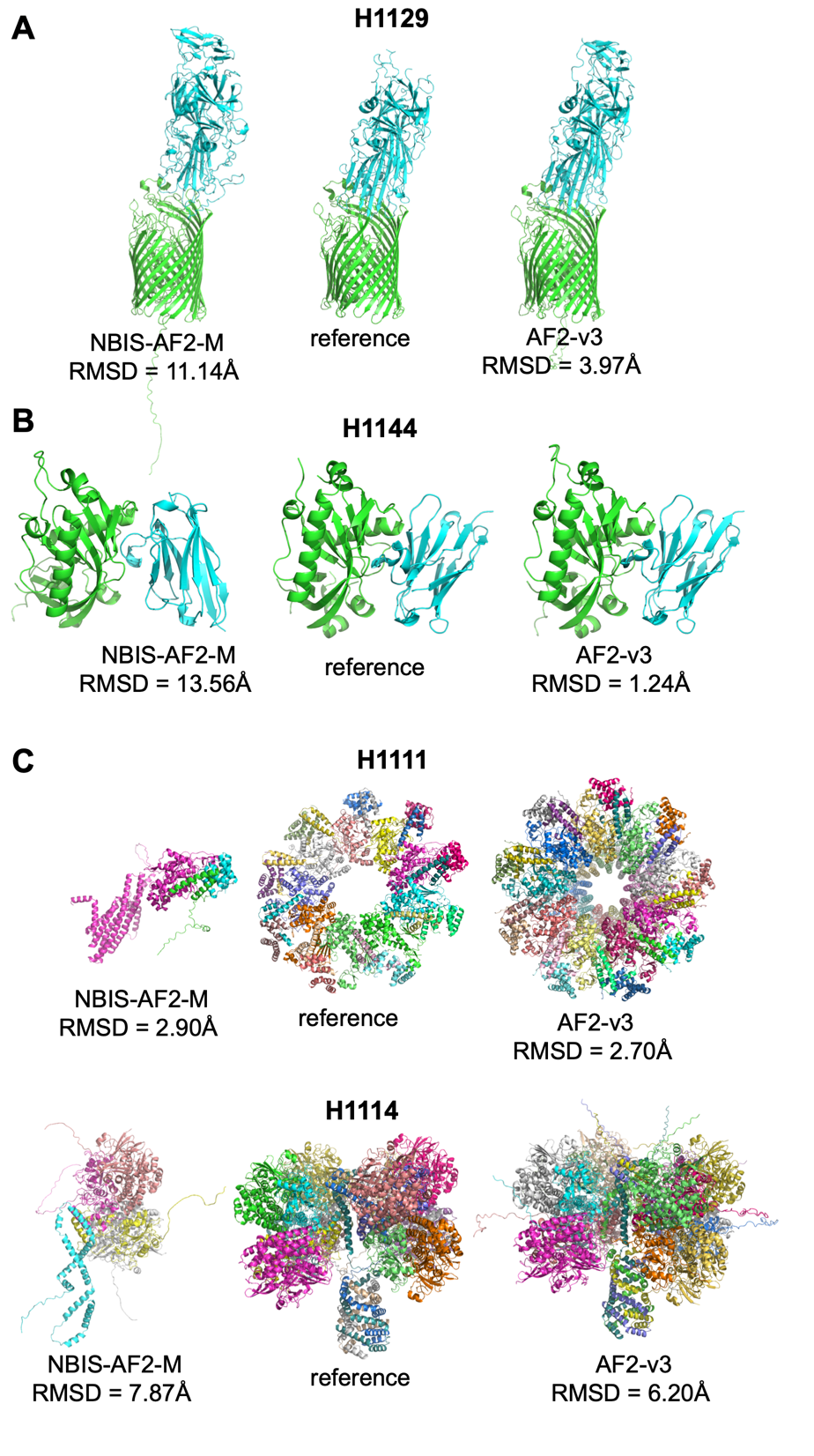

**Figure S11. AF2-v2.2 (NBIS-AF2M) and AF2-v2.3 model comparisons with and the reference structures.** All atom RMSDs are calculated with respect to the reference structure. **A.** H1129, **B.** H1144, **C.** H1111 and H1114. The RMSD of AF2-v2.3 models of H1111 and H1114 are based on the same asymmetric unit of NBIS-AF2-M models. The structures are colored according to their domains.

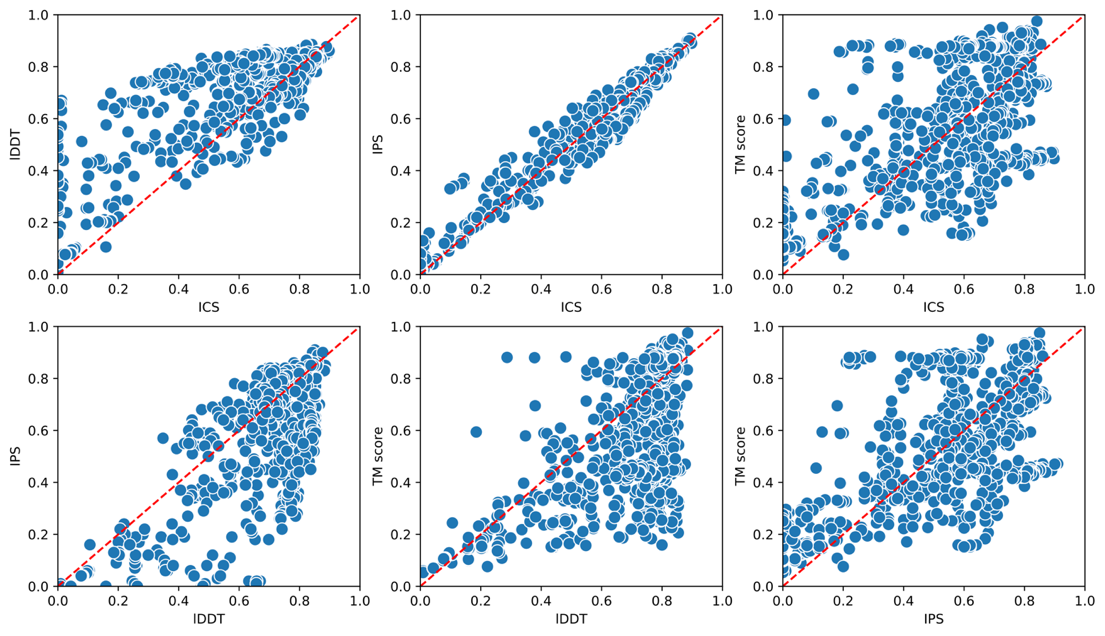

**Figure S12. The pairwise correlation of CASP15 metrics (ICS, IPS, lDDT, TM) by using all groups’ best models.** The TM score vs ICS plot reveals a "mismatch region" characterized by low TM values (<0.4) and high ICS values (>0.6). This region was observed in specific models from targets: H1114 (38 models), H1135 (3 models), H1137 (10 models), T1174 (8 models), and T1192 (2 models), which displayed a common issue of large interface dominance in the ICS score without accurate overall assembly prediction.

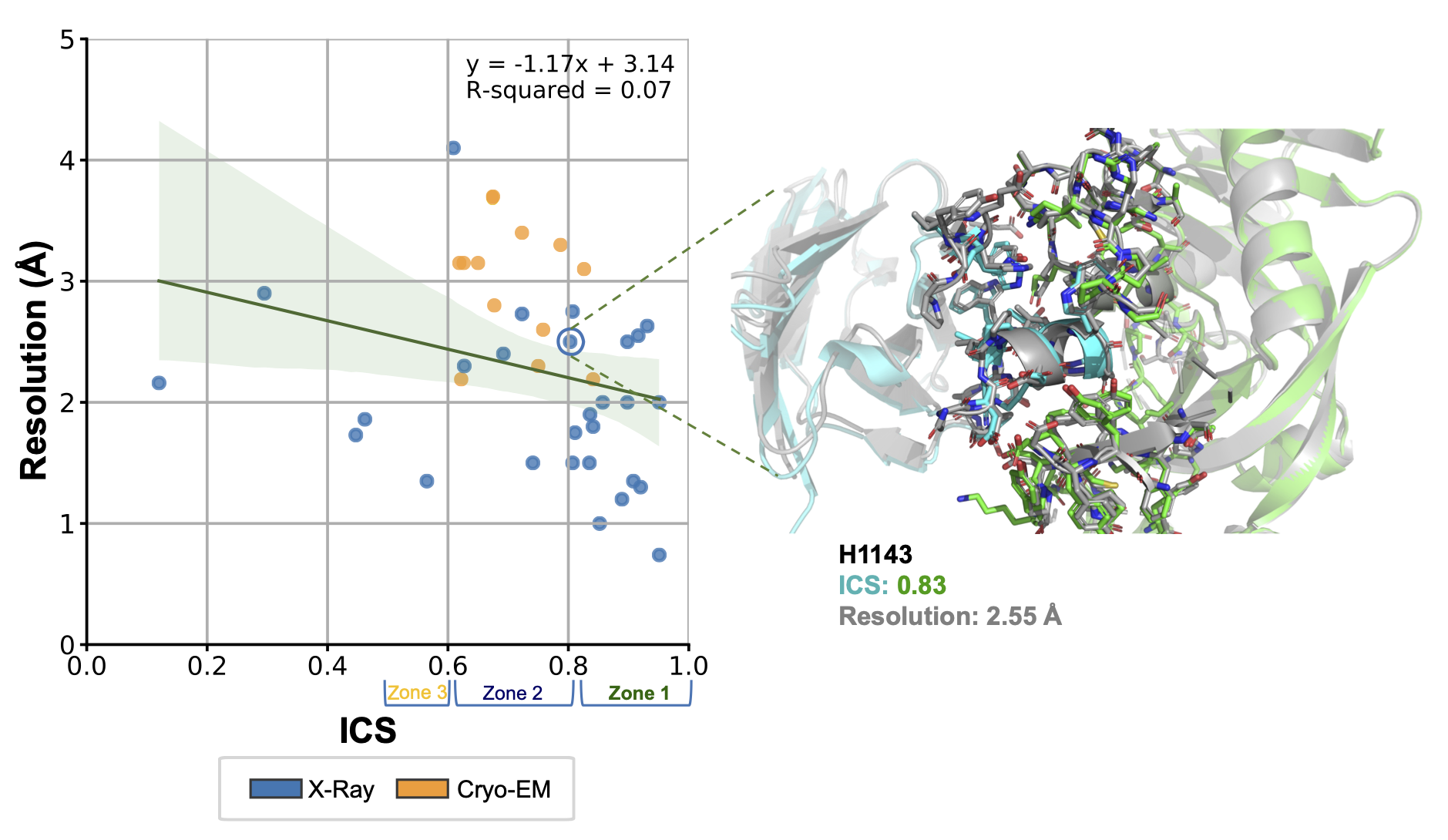

**Figure S13. The relationship between the resolution of the ground truth structure and the best ICS model generated for that target.** Zone1 denotes the ICS range of 1-0.8, while zone2 denotes the ICS range of 0.8-0.6 and zone3 denotes the ICS range 0.6-0.5. The inset highlights the interface prediction quality of a model of 0.8 ICS. Here, the interfacial side chains of the ground truth were shown in gray sticks, while the ones of the model were shown in green and cyan colors. The target and the best model come from the H1143 nanobody-target prediction challenge, where the ground truth structure has a resolution of 2.55 Å.

**TABLES:**

**Table S1. The closest templates** found for the subunits of easy and medium assembly cases, together with their target stoichiometries and prediction difficulties.

| Target ID | Stoichiometry | Difficulty | The closest template(s) found regarding different subunits | | | | | |
| --- | --- | --- | --- | --- | --- | --- | --- | --- |
|  |  |  | **A** | **B** | **C** | **D** | **H** | **I** |
| H1106 | A1B1 | easy | 7qii_b^1^ | 7qii_a^1^, 4ga2_a^2^ |  |  |  |  |
| H1111 | A9B9C9 | medium | 7alw |  |  |  |  |  |
| H1114 | A4B8C8 | medium |  | 2frv_c^3^ | 1h2a_a^4^ |  |  |  |
| H1114v1 | A4B2 | medium |  | 2frv_c^3^ |  |  |  |  |
| H1129 | A1B1 | medium | 3qlb_a^5^, 4cu4_a^6^ |  |  |  |  |  |
| H1134 | A1B1 | medium | 3ngm, 5gw8, 6mok_a^7^ | 6aow_a^8^ |  |  |  |  |
| H1135 | A9B3 | medium | 6r2i_a^9^ |  |  |  |  |  |
| H1137 | A1B1C1D1 E1F1G2H1I1 | medium |  |  |  |  | 7cge_a^10^ | 7cha_g^11^ |
| H1140 | A1B1 | medium | 4wfr^12^, 5ae0_a^13^ | 5iz0_a^14^ |  |  |  |  |
| H1141 | A1B1 | medium | 4wfr^12^, 5ae0_a^13^ | 5iz0_a^14^ |  |  |  |  |
| H1142 | A1B1 | medium | 4wfr^12^, 5ae0_a^13^ | 5iz0_a^14^ |  |  |  |  |
| H1143 | A1B1 | medium | 2ydd_a^15^, 4wfr^12^, 5ae0_a^13^ | 2ydd_ligand(2AM)^15^, 5iz0_a^14^ |  |  |  |  |
| H1144 | A1B1 | medium | 4wfr^12^, 5ae0_a^13^ | 5iz0_a^14^ |  |  |  |  |
| H1151 | A1B1 | easy | 7kug_d^16^ | 7kug_c^16^ |  |  |  |  |
| H1157 | A1B1 | medium | 2ri9_b^17^ | 5xf7_a^18^ |  |  |  |  |
| H1166v1 | A1B1 (H+L=1chain) | medium | 2vxv^19^ | 7dyd_a^20^ |  |  |  |  |
| H1167v1 | A1B1 (H+L=1chain) | medium | 2vxv^19^ | 7dyd_a^20^ |  |  |  |  |
| H1168v1 | A1B1 (H+L=1chain) | medium | 7r98_d^21^, 2vxv^19^ | 7r98_a^21^, 7dyd_a^20^ |  |  |  |  |
| H1171 | A6B1 | easy | 1ixs^22^, 3pfi | 1ixs^22^, 7oa5^23^ |  |  |  |  |
| H1172 | A6B2 | easy | 1ixs^22^, 3pfi | 1ixs^22^, 7oa5^23^ |  |  |  |  |
| H1185 | A1B1C1D1 | easy | 7ej6^24^ | 7ej6^24^ | 5fos^25^, 7ej6^24^ | 5fos^25^, 7ej6^24^ |  |  |
| T1109o | A2 | easy | 6unf^26^ |  |  |  |  |  |
| T1110o | A2 | easy | 6unf^26^ |  |  |  |  |  |
| T1115o | A16 | medium | 7vhq^27^ |  |  |  |  |  |
| T1121o | A2 | medium | 2q2e_a^28^, 1d3y_a^29^ |  |  |  |  |  |
| T1123o | A2 | medium | 3qsq_a^30^ |  |  |  |  |  |
| T1124o | A2 | easy | 2r3s |  |  |  |  |  |
| T1127o | A2 | easy | 2fe7 |  |  |  |  |  |
| T1132o | A6 | medium | 2bbe_a, 2hiq_a |  |  |  |  |  |
| T1153o | A2 | medium | 5uvg^31^ |  |  |  |  |  |
| T1160o | A2 | easy | 7dxx^32^, 7dxy^32^ |  |  |  |  |  |
| T1161o | A2 | easy | 7dxx^32^, 7dxy^32^ |  |  |  |  |  |
| T1170o | A6 | easy | 3pfi |  |  |  |  |  |
| T1178o | A2 | easy | 5ewo^33^, 5w1n^34^ |  |  |  |  |  |
| T1179o | A2 | easy | 5w1n^34^ |  |  |  |  |  |
| T1181o | A3 | easy | 6tgf^35^ |  |  |  |  |  |
| T1192o | A10 | easy | 5jrb^36^ |  |  |  |  |  |

**Table S2.** **The ranking of CASP15 assembly groups**, according to mean scores of their first submitted models over 41 targets. The servers are marked with *.

| Number | Group name | Group no | Mean of sum of Z-scores (>0.0) | Standard deviation | Rank sum z-score (>0.0) |
| --- | --- | --- | --- | --- | --- |
| 1 | Zheng | 374 | 34.44 | 0.71 | 1 |
| 2 | Venclovas | 494 | 28.43 | 0.68 | 2 |
| 3 | Wallner | 36 | 27.45 | 0.94 | 3 |
| 4 | Yang-Multimer* | 239 | 24.09 | 0.62 | 4 |
| 5 | Yang | 439 | 23.58 | 0.61 | 5 |
| 6 | Kiharalab | 119 | 21.29 | 0.49 | 6 |
| 7 | MULTICOM_human | 3 | 20.21 | 0.32 | 7 |
| 8 | Manifold | 248 | 19.79 | 0.49 | 8 |
| 9 | McGuffin | 180 | 19.41 | 0.47 | 9 |
| 10 | MULTICOM | 367 | 19.17 | 0.33 | 10 |
| 11 | Manifold-E* | 35 | 18.40 | 0.49 | 11 |
| 12 | MULTICOM_qa* | 86 | 17.91 | 0.30 | 12 |
| 13 | PEZYFoldings | 278 | 17.51 | 0.50 | 13 |
| 14 | DFolding-server* | 288 | 16.60 | 0.54 | 14 |
| 15 | MULTICOM_deep* | 158 | 15.89 | 0.28 | 15 |
| 16 | CoDock | 444 | 15.87 | 0.57 | 16 |
| 17 | BAKER | 185 | 15.52 | 0.45 | 17 |
| 18 | UltraFold | 54 | 15.39 | 0.46 | 18 |
| 19 | BeijingAIProtein | 399 | 15.33 | 0.46 | 19 |
| 20 | UltraFold_Server* | 125 | 15.32 | 0.46 | 20 |
| 21 | Elofsson | 320 | 15.30 | 0.25 | 21 |
| 22 | Takeda-Shitaka_Lab | 348 | 15.25 | 0.30 | 22 |
| 23 | MultiFOLD* | 462 | 14.86 | 0.45 | 23 |
| 24 | MUFold_H | 360 | 14.73 | 0.32 | 24 |
| 25 | colabfold_human | 461 | 13.99 | 0.34 | 25 |
| 26 | MUFold* | 298 | 13.75 | 0.26 | 26 |
| 27 | Pierce | 314 | 13.70 | 0.60 | 27 |
| 28 | Kiharalab_Server* | 131 | 13.19 | 0.56 | 28 |
| 29 | ColabFold* | 446 | 12.46 | 0.31 | 29 |
| 30 | NBIS-AF2-multimer* | 390 | 11.97 | 0.24 | 30 |
| 31 | RaptorX-Multimer* | 71 | 11.63 | 0.27 | 31 |
| 32 | Grudinin | 150 | 11.60 | 0.24 | 32 |
| 33 | DMP | 477 | 11.14 | 0.42 | 33 |
| 34 | Yang-Server* | 229 | 10.24 | 0.37 | 34 |
| 35 | SHT | 147 | 9.95 | 0.36 | 35 |
| 36 | DFolding-refine* | 73 | 9.10 | 0.45 | 36 |
| 37 | ShanghaiTech | 225 | 9.05 | 0.47 | 37 |
| 38 | ClusPro | 350 | 8.82 | 0.23 | 38 |
| 39 | GuijunLab-Human | 169 | 8.65 | 0.22 | 39 |
| 40 | Coqualia | 434 | 8.45 | 0.36 | 40 |
| 41 | GuijunLab-Assembly* | 98 | 8.39 | 0.26 | 41 |
| 42 | FTBiot0119 | 165 | 7.48 | 0.31 | 42 |
| 43 | Shen-CAPRI | 493 | 7.40 | 0.49 | 43 |
| 44 | trComplex | 423 | 7.26 | 0.25 | 44 |
| 45 | GinobiFold-SER* | 11 | 6.61 | 0.33 | 45 |
| 46 | GuijunLab-DeepDA* | 188 | 6.29 | 0.20 | 46 |
| 47 | TRFold | 187 | 5.98 | 0.23 | 47 |
| 48 | Zou | 205 | 5.97 | 0.19 | 48 |
| 49 | GinobiFold | 227 | 5.41 | 0.22 | 49 |
| 50 | Manifold-X | 304 | 5.31 | 0.29 | 50 |
| 51 | WL_team | 257 | 5.18 | 0.22 | 51 |
| 52 | DELCLAB | 447 | 5.14 | 0.43 | 52 |
| 53 | Agemo_mix | 92 | 4.99 | 0.23 | 53 |
| 54 | Kozakov-Vajda | 291 | 4.90 | 0.19 | 54 |
| 55 | UNRES | 91 | 4.77 | 0.19 | 55 |
| 56 | DFolding | 74 | 4.73 | 0.29 | 56 |
| 57 | ShanghaiTech-TS-SER* | 133 | 4.61 | 0.21 | 57 |
| 58 | FoldEver-Hybrid | 385 | 4.52 | 0.14 | 58 |
| 59 | FoldEver* | 245 | 4.21 | 0.16 | 59 |
| 60 | Manifold-LC-E* | 46 | 3.95 | 0.25 | 60 |
| 61 | bio3d | 397 | 3.74 | 0.36 | 61 |
| 62 | Fernandez-Recio | 312 | 3.72 | 0.17 | 62 |
| 63 | ChaePred | 398 | 3.38 | 0.14 | 63 |
| 64 | OpenFold | 441 | 2.76 | 0.24 | 64 |
| 65 | OpenFold-SingleSeq | 433 | 2.76 | 0.24 | 64 |
| 66 | AIchemy_LIG3 | 347 | 2.67 | 0.16 | 66 |
| 67 | AIchemy_LIG2 | 456 | 2.64 | 0.16 | 67 |
| 68 | AIchemy_LIG | 325 | 2.64 | 0.16 | 67 |
| 69 | ddquest | 472 | 2.21 | 0.20 | 68 |
| 70 | KORP-PL | 352 | 2.14 | 0.18 | 69 |
| 71 | Convex-PL-R | 460 | 1.98 | 0.18 | 70 |
| 72 | zax | 122 | 1.95 | 0.14 | 71 |
| 73 | UTMB | 201 | 1.87 | 0.17 | 72 |
| 74 | Convex-PL | 338 | 1.67 | 0.15 | 73 |
| 75 | TB_model_prediction | 199 | 1.28 | 0.11 | 74 |
| 76 | XRC_VU* | 215 | 1.12 | 0.13 | 75 |
| 77 | Graphen_Medical | 97 | 0.91 | 0.06 | 76 |
| 78 | ESM-single-sequence | 67 | 0.87 | 0.10 | 77 |
| 79 | Cerebra* | 315 | 0.52 | 0.06 | 78 |
| 80 | Gonglab-THU | 52 | 0.52 | 0.06 | 78 |
| 81 | noxelis | 236 | 0.40 | 0.06 | 80 |
| 82 | TensorLab | 132 | 0.35 | 0.04 | 81 |
| 83 | Panlab | 234 | 0.34 | 0.04 | 82 |
| 84 | GuijunLab-Meta* | 481 | 0.33 | 0.05 | 83 |
| 85 | wuqi* | 370 | 0.00 | 0.00 | 84 |
| 86 | FALCON2* | 368 | 0.00 | 0.00 | 84 |
| 87 | FALCON0* | 333 | 0.00 | 0.00 | 84 |

**Table S3. The ranking of CASP15 interdomain groups**, according to their first submitted models over 24 evaluation units.

| Number | Group name | Group no | Number of attended evaluation units | Sum z-score (>0.0) | Rank sum z-score (>0.0) |
| --- | --- | --- | --- | --- | --- |
| 1 | UM-TBM | 162 | 20 | 35.5277 | 1 |
| 2 | Yang-Server | 229 | 20 | 24.9602 | 2 |
| 3 | Yang | 439 | 20 | 19.7115 | 3 |
| 4 | PEZYFoldings | 278 | 15 | 18.0578 | 4 |
| 5 | Manifold | 248 | 19 | 14.9308 | 5 |
| 6 | Venclovas | 494 | 19 | 14.5386 | 6 |
| 7 | server_124 | 383 | 20 | 14.081 | 7 |
| 8 | DFolding | 074 | 20 | 13.1098 | 8 |
| 9 | bench | 008 | 20 | 12.0811 | 9 |
| 10 | BAKER-SERVER | 443 | 20 | 12.003 | 10 |
| 11 | Manifold-E | 035 | 19 | 11.6732 | 11 |
| 12 | DFolding-server | 288 | 19 | 11.587 | 12 |
| 13 | server_126 | 403 | 20 | 10.9291 | 13 |
| 14 | Shennong | 466 | 20 | 10.1011 | 14 |
| 15 | RaptorX | 166 | 20 | 9.3845 | 15 |
| 16 | IntFOLD7 | 151 | 19 | 9.0429 | 16 |
| 17 | BAKER | 185 | 20 | 8.862 | 17 |
| 18 | server_123 | 018 | 20 | 8.5973 | 18 |
| 19 | Asclepius | 204 | 19 | 8.5891 | 19 |
| 20 | MultiFOLD | 462 | 18 | 8.2488 | 20 |
| 21 | DFolding-refine | 073 | 19 | 8.1694 | 21 |
| 22 | B11L | 208 | 18 | 8.0841 | 22 |
| 23 | DMP | 477 | 14 | 7.7417 | 23 |
| 24 | MUFold_H | 360 | 20 | 7.5603 | 24 |
| 25 | hFold | 353 | 16 | 7.1212 | 25 |
| 26 | OpenFold-SingleSeq | 433 | 19 | 7.0733 | 26 |
| 27 | OpenFold | 441 | 19 | 7.0733 | 26 |
| 28 | ShanghaiTech | 225 | 20 | 7.0584 | 28 |
| 29 | ManiFold-serv | 450 | 19 | 6.9819 | 29 |
| 30 | Graphen_Medical | 097 | 14 | 6.8295 | 30 |
| 31 | AP_1 | 269 | 20 | 6.8095 | 31 |
| 32 | Elofsson | 320 | 19 | 6.6876 | 32 |
| 33 | Agemo_mix | 092 | 19 | 6.6477 | 33 |
| 34 | Panlab | 234 | 20 | 6.5871 | 34 |
| 35 | McGuffin | 180 | 19 | 6.5509 | 35 |
| 36 | TRFold | 187 | 17 | 6.4502 | 36 |
| 37 | MULTICOM_refine | 475 | 19 | 6.421 | 37 |
| 38 | server_122 | 261 | 20 | 6.1989 | 38 |
| 39 | BeijingAIProtein | 399 | 17 | 6.1858 | 39 |
| 40 | UltraFold | 054 | 17 | 6.1858 | 39 |
| 41 | UltraFold_Server | 125 | 18 | 6.1858 | 39 |
| 42 | server_125 | 264 | 20 | 6.0672 | 42 |
| 43 | Agemo | 478 | 17 | 5.9827 | 43 |
| 44 | FTBiot0119 | 165 | 20 | 5.9501 | 44 |
| 45 | ChaePred | 398 | 20 | 5.7616 | 45 |
| 46 | WL_team | 257 | 19 | 5.7417 | 46 |
| 47 | Kiharalab_Server | 131 | 20 | 5.7358 | 47 |
| 48 | trComplex | 423 | 17 | 5.6135 | 48 |
| 49 | hFold_human | 342 | 18 | 5.5688 | 49 |
| 50 | Kiharalab | 119 | 20 | 5.494 | 50 |
| 51 | MULTICOM_deep | 158 | 19 | 5.4897 | 51 |
| 52 | Seder2022easy | 455 | 19 | 5.3198 | 52 |
| 53 | GuijunLab-DeepDA | 188 | 19 | 4.8945 | 53 |
| 54 | XRC_VU | 215 | 7 | 4.8243 | 54 |
| 55 | ColabFold | 446 | 20 | 4.7985 | 55 |
| 56 | colabfold_human | 461 | 20 | 4.7985 | 55 |
| 57 | GuijunLab-Assembly | 098 | 19 | 4.3734 | 57 |
| 58 | Wallner | 037 | 18 | 4.1617 | 58 |
| 59 | FoldEver | 245 | 19 | 3.9173 | 59 |
| 60 | MULTICOM | 367 | 18 | 3.9011 | 60 |
| 61 | MULTICOM_human | 003 | 18 | 3.6862 | 61 |
| 62 | GuijunLab-Meta | 481 | 19 | 3.6577 | 62 |
| 63 | MULTICOM_qa | 086 | 20 | 3.5934 | 63 |
| 64 | GuijunLab-Human | 169 | 19 | 3.4694 | 64 |
| 65 | FoldEver-Hybrid | 385 | 14 | 3.3126 | 65 |
| 66 | MULTICOM_egnn | 120 | 20 | 3.1465 | 66 |
| 67 | GinobiFold | 227 | 17 | 2.7459 | 67 |
| 68 | Coqualia | 434 | 17 | 2.7459 | 67 |
| 69 | Cerebra | 315 | 17 | 2.5507 | 69 |
| 70 | MUFold | 298 | 20 | 2.4377 | 70 |
| 71 | GuijunLab-RocketX | 089 | 19 | 2.424 | 71 |
| 72 | GuijunLab-Threader | 282 | 18 | 2.4147 | 72 |
| 73 | Bhattacharya | 275 | 19 | 2.372 | 73 |
| 74 | SHT | 147 | 20 | 2.2983 | 74 |
| 75 | BhageerathH-Pro | 212 | 14 | 2.2781 | 75 |
| 76 | GinobiFold-SER | 011 | 16 | 2.2577 | 76 |
| 77 | FALCON2 | 368 | 20 | 2.2265 | 77 |
| 78 | FALCON0 | 333 | 20 | 2.2265 | 77 |
| 79 | hks1988 | 354 | 20 | 2.1535 | 79 |
| 80 | NBIS-AF2-standard | 270 | 20 | 2.1141 | 80 |
| 81 | Pan_Server | 219 | 19 | 2.0335 | 81 |
| 82 | Gonglab-THU | 052 | 17 | 1.8164 | 82 |
| 83 | DELCLAB | 447 | 18 | 1.1885 | 83 |
| 84 | ESM-single-sequence | 067 | 9 | 1.1811 | 84 |
| 85 | UNRES | 091 | 14 | 1.1657 | 85 |
| 86 | QUIC | 117 | 20 | 1.1187 | 86 |
| 87 | PICNIC | 276 | 20 | 1.1002 | 87 |
| 88 | ShanghaiTech-TS-SER | 133 | 16 | 0.8613 | 88 |
| 89 | Seder2022hard | 216 | 10 | 0.591 | 89 |
| 90 | SHORTLE | 064 | 1 | 0.59 | 90 |
| 91 | wuqi | 370 | 9 | 0.4176 | 91 |
| 92 | MESHI_server | 427 | 2 | 0.1894 | 92 |
| 93 | EMBER3D | 140 | 10 | 0.1591 | 93 |
| 94 | Manifold-LC-E | 046 | 1 | 0.0809 | 94 |
| 95 | Manifold-X | 304 | 1 | 0.0809 | 94 |
| 96 | RostlabUeFOFold | 123 | 4 | 0.0346 | 96 |
| 97 | MESHI | 362 | 1 | 0 | 97 |
| 98 | ACOMPMOD | 280 | 1 | 0 | 97 |

**Table S4. The CASP15 interdomain targets** and the top performing predictor for that target (ranked according to sum of z-scores>0).

| Target ID | First ranking group |
| --- | --- |
| T1125 | UM-TBM |
| T1125-D12 | UM-TBM |
| T1125-D23 | UM-TBM |
| T1125-D34 | UM-TBM |
| T1125-D45 | PEZYFoldings |
| T1125-D56 | BAKER-SERVER |
| T1145 | MULTICOM_deep |
| T1154 | Yang-Server |
| T1158 | MUFold_H |
| T1165 | Yang |
| T1165-D12 | PEZYFoldings |
| T1165-D16 | UltraFold |
| T1165-D23 | Agemo |
| T1165-D36 | Yang |
| T1165-D56 | OpenFold-SingleSeq |
| T1169 | Yang-Server |
| T1169-D12 | Yang-Server |
| T1169-D23 | PEZYFoldings |
| T1169-D34 | Venclovas |
| T1180 | OpenFold-SingleSeq |

**REFERENCES:**

1.	Wu K, Moore JA, Miller MD, et al. Expanding the eukaryotic genetic code with a biosynthesized 21st amino acid. *Protein Science*. 2022;31(10). doi:10.1002/pro.4443

2.	Kassube SA, Stuwe T, Lin DH, et al. Crystal Structure of the N-Terminal Domain of Nup358/RanBP2. *J Mol Biol*. 2012;423(5):752-765. doi:10.1016/j.jmb.2012.08.026

3.	Volbeda A, Garcin E, Piras C, et al. Structure of the [NiFe] Hydrogenase Active Site:  Evidence for Biologically Uncommon Fe Ligands. *J Am Chem Soc*. 1996;118(51):12989-12996. doi:10.1021/ja962270g

4.	Higuchi Y, Yagi T, Yasuoka N. Unusual ligand structure in Ni–Fe active center and an additional Mg site in hydrogenase revealed by high resolution X-ray structure analysis. *Structure*. 1997;5(12):1671-1680. doi:10.1016/S0969-2126(97)00313-4

5.	Brillet K, Reimmann C, Mislin GLA, et al. Pyochelin Enantiomers and Their Outer-Membrane Siderophore Transporters in Fluorescent Pseudomonads: Structural Bases for Unique Enantiospecific Recognition. *J Am Chem Soc*. 2011;133(41):16503-16509. doi:10.1021/ja205504z

6.	Mathavan I, Zirah S, Mehmood S, et al. Structural basis for hijacking siderophore receptors by antimicrobial lasso peptides. *Nat Chem Biol*. 2014;10(5):340-342. doi:10.1038/nchembio.1499

7.	Mohan K, Ueda G, Kim AR, et al. Topological control of cytokine receptor signaling induces differential effects in hematopoiesis. *Science (1979)*. 2019;364(6442). doi:10.1126/science.aav7532

8.	Kalas V, Hibbing ME, Maddirala AR, et al. Structure-based discovery of glycomimetic FmlH ligands as inhibitors of bacterial adhesion during urinary tract infection. *Proceedings of the National Academy of Sciences*. 2018;115(12). doi:10.1073/pnas.1720140115

9.	Gurusaran M, Davies OR. A molecular mechanism for LINC complex branching by structurally diverse SUN-KASH 6:6 assemblies. *Elife*. 2021;10. doi:10.7554/eLife.60175

10.	Chi X, Fan Q, Zhang Y, et al. Structural mechanism of phospholipids translocation by MlaFEDB complex. *Cell Res*. 2020;30(12):1127-1135. doi:10.1038/s41422-020-00404-6

11.	Zhou C, Shi H, Zhang M, et al. Structural Insight into Phospholipid Transport by the MlaFEBD Complex from P. aeruginosa. *J Mol Biol*. 2021;433(13):166986. doi:10.1016/j.jmb.2021.166986

12.	Raasakka A, Myllykoski M, Laulumaa S, et al. Determinants of ligand binding and catalytic activity in the myelin enzyme 2′,3′-cyclic nucleotide 3′-phosphodiesterase. *Sci Rep*. 2015;5(1):16520. doi:10.1038/srep16520

13.	Raasakka A, Myllykoski M, Laulumaa S, et al. Determinants of ligand binding and catalytic activity in the myelin enzyme 2′,3′-cyclic nucleotide 3′-phosphodiesterase. *Sci Rep*. 2015;5(1):16520. doi:10.1038/srep16520

14.	Marcotte DJ, Liu Y, Little K, et al. Structural determinant for inducing RORgamma specific inverse agonism triggered by a synthetic benzoxazinone ligand. *BMC Struct Biol*. 2016;16(1):7. doi:10.1186/s12900-016-0059-3

15.	Myllykoski M, Raasakka A, Han H, Kursula P. Myelin 2′,3′-Cyclic Nucleotide 3′-Phosphodiesterase: Active-Site Ligand Binding and Molecular Conformation. *PLoS One*. 2012;7(2):e32336. doi:10.1371/journal.pone.0032336

16.	Wan T, Horová M, Beltran DG, Li S, Wong HX, Zhang LM. Structural insights into the functional divergence of WhiB-like proteins in Mycobacterium tuberculosis. *Mol Cell*. 2021;81(14):2887-2900.e5. doi:10.1016/j.molcel.2021.06.002

17.	Lobsanov YD, Yoshida T, Desmet T, et al. Modulation of activity by Arg407: structure of a fungal α-1,2-mannosidase in complex with a substrate analogue. *Acta Crystallogr D Biol Crystallogr*. 2008;64(3):227-236. doi:10.1107/S0907444907065572

18.	Li H, Yang K, Wang W, et al. Crystal and solution structures of human protein-disulfide isomerase-like protein of the testis (PDILT) provide insight into its chaperone activity. *Journal of Biological Chemistry*. 2018;293(4):1192-1202. doi:10.1074/jbc.M117.797290

19.	Argiriadi MA, Xiang T, Wu C, Ghayur T, Borhani DW. Unusual Water-mediated Antigenic Recognition of the Proinflammatory Cytokine Interleukin-18. *Journal of Biological Chemistry*. 2009;284(36):24478-24489. doi:10.1074/jbc.M109.023887

20.	Hsu JN, Chen JS, Lin SM, et al. Targeting the N-Terminus Domain of the Coronavirus Nucleocapsid Protein Induces Abnormal Oligomerization via Allosteric Modulation. *Front Mol Biosci*. 2022;9. doi:10.3389/fmolb.2022.871499

21.	Ye Q, Lu S, Corbett KD. Structural Basis for SARS-CoV-2 Nucleocapsid Protein Recognition by Single-Domain Antibodies. *Front Immunol*. 2021;12. doi:10.3389/fimmu.2021.719037

22.	Yamada K, Miyata T, Tsuchiya D, et al. Crystal Structure of the RuvA-RuvB Complex. *Mol Cell*. 2002;10(3):671-681. doi:10.1016/S1097-2765(02)00641-X

23.	Roe SM, Barlow T, Brown T, et al. Crystal Structure of an Octameric RuvA–Holliday Junction Complex. *Mol Cell*. 1998;2(3):361-372. doi:10.1016/S1097-2765(00)80280-4

24.	Xu J, Zhao L, Peng S, et al. Mechanisms of distinctive mismatch tolerance between Rad51 and Dmc1 in homologous recombination. *Nucleic Acids Res*. 2021;49(22):13135-13149. doi:10.1093/nar/gkab1141

25.	Moschetti T, Sharpe T, Fischer G, et al. Engineering Archeal Surrogate Systems for the Development of Protein–Protein Interaction Inhibitors against Human RAD51. *J Mol Biol*. 2016;428(23):4589-4607. doi:10.1016/j.jmb.2016.10.009

26.	Dasgupta M, Budday D, de Oliveira SHP, et al. Mix-and-inject XFEL crystallography reveals gated conformational dynamics during enzyme catalysis. *Proceedings of the National Academy of Sciences*. 2019;116(51):25634-25640. doi:10.1073/pnas.1901864116

27.	Ma C, Wang C, Luo D, et al. Structural insights into the membrane microdomain organization by SPFH family proteins. *Cell Res*. 2022;32(2):176-189. doi:10.1038/s41422-021-00598-3

28.	Corbett KD, Benedetti P, Berger JM. Holoenzyme assembly and ATP-mediated conformational dynamics of topoisomerase VI. *Nat Struct Mol Biol*. 2007;14(7):611-619. doi:10.1038/nsmb1264

29.	Nichols MD. Structure and function of an archaeal topoisomerase VI subunit with homology to the meiotic recombination factor Spo11. *EMBO J*. 1999;18(21):6177-6188. doi:10.1093/emboj/18.21.6177

30.	Dong J, Dong L, Méndez E, Tao Y. Crystal structure of the human astrovirus capsid spike. *Proceedings of the National Academy of Sciences*. 2011;108(31):12681-12686. doi:10.1073/pnas.1104834108

31.	Airola M V., Shanbhogue P, Shamseddine AA, et al. Structure of human nSMase2 reveals an interdomain allosteric activation mechanism for ceramide generation. *Proceedings of the National Academy of Sciences*. 2017;114(28). doi:10.1073/pnas.1705134114

32.	Yagi S, Padhi AK, Vucinic J, et al. Seven Amino Acid Types Suffice to Create the Core Fold of RNA Polymerase. *J Am Chem Soc*. 2021;143(39):15998-16006. doi:10.1021/jacs.1c05367

33.	York RL, Yousefi PA, Bogdanoff W, Haile S, Tripathi S, DuBois RM. Structural, Mechanistic, and Antigenic Characterization of the Human Astrovirus Capsid. *J Virol*. 2016;90(5):2254-2263. doi:10.1128/JVI.02666-15

34.	Bogdanoff WA, Perez EI, López T, Arias CF, DuBois RM. Structural Basis for Escape of Human Astrovirus from Antibody Neutralization: Broad Implications for Rational Vaccine Design. *J Virol*. 2018;92(1). doi:10.1128/JVI.01546-17

35.	Irmscher T, Roske Y, Gayk I, et al. Pantoea stewartii WceF is a glycan biofilm-modifying enzyme with a bacteriophage tailspike-like fold. *Journal of Biological Chemistry*. 2021;296:100286. doi:10.1016/j.jbc.2021.100286

36.	Saotome M, Saito K, Onodera K, Kurumizaka H, Kagawa W. Structure of the human DNA-repair protein RAD52 containing surface mutations. *Acta Crystallogr F Struct Biol Commun*. 2016;72(8):598-603. doi:10.1107/S2053230X1601027X
